# A Mixed Effect Similarity Matrix Regression Model (SMRmix) for Integrating Multiple Microbiome Datasets at Community Level

**DOI:** 10.1101/2024.03.10.584315

**Authors:** Mengyu He, Ni Zhao

## Abstract

**Background:** Recent studies have highlighted the importance of human microbiota in our health and diseases. However, in many areas of research, individual microbiome studies often offer inconsistent results due to the limited sample sizes and the heterogeneity in study populations and experimental procedures. This inconsistency underscores the necessity for integrative analysis of multiple microbiome datasets. Despite the critical need, statistical methods that incorporate multiple microbiome datasets and account for the study heterogeneity are not available in the literature.

**Methods:** In this paper, we develop a mixed effect similarity matrix regression (SMRmix) approach for identifying community level microbiome shifts between outcomes. SMRmix has a close connection with the microbiome kernel association test, one of the most popular approaches for such a task but is only applicable when we have a single study. SMRmix enables researchers to consolidate findings from diverse microbiome studies.

**Results:** Via extensive simulations, we show that SMRmix has well-controlled type I error and higher power than some potential competitors. We applied the SMRmix to two real-world datasets. The first, from the HIV-reanalysis consortium, integrated data from 17 studies on gut dysbiosis in HIV. Our analysis confirmed consistent associations between the gut microbiome and HIV infection as well as MSM (men who have sex with men) status, demonstrating greater power than competing methods. The second dataset involved 11 studies on the gut microbiome in colorectal cancer; analysis with SMRmix confirmed significant dysbiosis in affected individuals compared to healthy controls.

**Conclusion:** The development of SMRmix enables the integration of multiple studies and effectively managing study heterogeneity, and provides a powerful tool for uncovering consistent associations between diseases and community-level microbiome data.

## 1 Introduction

Microbiome research has rapidly expanded in recent years, revealing complex relationships between microbial communities and human health. However, individual studies that explore the relationship between the microbiome and disease frequently report inconsistent results. For example, studies on the gut microbiome and HIV infections have shown conflicting results. While most studies found a decreased microbial alpha diversity in HIV-infected individuals compared to uninfected individuals [1, 2, 3, 4, 5, 6, 7, 8, 9, 10, 11], some studies detected no significant difference [12, 13, 14, 15] and even an increase of microbial diversity [16]. Research on structural differences in microbial communities (e.g., beta diversity) between HIV-positive and HIV-negative individuals has also yielded mixed results. Some studies report significant differences in the microbiome beta diversity of HIV-infected versus uninfected individuals [2, 14], while others suggest that these differences may be attributed more to variations in sexual behavior, especially MSM (men who have sex with men) status, rather than HIV infection itself [3]. Similar inconsistencies in microbial diversity and community structure have also been observed in other diseases, highlighting a broader challenge in microbiome research. These inconsistencies can arise from small sample sizes, population heterogeneity, varied study designs, and differences in analytical techniques, which may lead to bias and limit the generalizability of findings.

To address these issues and enhance reproducibility, integrative analysis of multiple microbiome datasets is essential to aggregate information across multiple studies. In this paper, we focus on the association test between the overall microbiome community profile, measured by beta-diversities, and outcomes of interest while incorporating multiple datasets. For such tasks within a single study, there are two popular approaches: distance-based analysis [17, 18] and the microbiome regression kernel association test (MiRKAT) [19]. The distance-based approach involves computing a beta-diversity distance between individuals’ microbiome profiles and testing for association with the outcome of interest via permutation. MiRKAT utilizes a semi-parametric kernel machine regression in which a kernel similarity matrix, derived from the beta-diversity matrix, is assessed for association with the outcome. Here, the kernel similarity matrix is transformed from of a beta-diversity matrix. Both methods require an assessment of similarity or dissimilarity(distance) between all pairs of samples.

When extending to multiple studies, neither distance-based methods nor MiRKAT are directly applicable due to the lack of a complete pairwise distance matrix across all subjects and studies. In cases where data are obtained from public resources without access to original sequencing files, differences in preprocessing steps across studies make it impossible to construct consistent betadiversity matrices—distinct OTU (operational taxonomic unit) assignments and phylogenetic trees are common. Even when re-processing data across studies is feasible, differential biases introduced through various sequencing protocols can invalidate beta diversity measurements between samples from different studies. Thus, in most cases, only within-study distances are available, while between-study distances are missing.

Herein, we propose a novel statistical method, SMRmix, to test the association between microbiome communities and outcomes of interests by aggregating information across multiple studies. Our method is built upon similarity matrix regression (SMR) and only requires within-study distances. Instead of modeling the direct relationship between traits and microbiome similarity, SMR regresses pairwise trait similarity on pairwise microbiome similarity for association testing. Trait similarity is calculated as the outer product of residuals from a null regression model, and microbiome similarity is calculated from the within-study beta diversity matrix. Associations are then detected using a score test. We extend SMR by incorporating mixed effects, vectorizing similarity matrices, and deriving the corresponding score test statistic through the likelihood. The new method, SMRmix, enables us to account for study-level random effects and avoid reliance on inaccessible between-study distances. Our method also enables development of an “optimal” omnibus test, utilizing multiple distances within a single study to enhance power.

Through extensive simulations, we demonstrate that the proposed SMRmix has well-controlled type I error rates, and higher or comparable power than potential competitors, such as CSKAT [20] and GLMM-MiRKAT [21], both of which require complete distance matrices across all studies, which are often inaccessible. We apply SMRmix to two datasets: (1) the HIV-reanalysis consortium dataset, comprising 17 studies on gut dysbiosis in HIV, where we observed strong associations between gut microbiome composition and both HIV infection and MSM (men who have sex with men) status, and (2) a dataset of 11 studies on gut microbiome in colorectal cancer (CRC), confirming significant dysbiosis in affected individuals compared to healthy controls. These analyses illustrate the capability of SMRmix to detect consistent patterns of microbiome composition composition across different conditions.

## 2 Materials and methods

SMRmix evaluates the association between a binary or continuous outcome and microbiome communities by integrating data from multiple microbiome studies. It extends similarity matrix regression to incorporate multiple studies, utilizing only intra-study kernel matrices, thereby eliminating the need for between-study similarity assessments. In what follows, we first present the microbiome regression-based kernel association test for a single study and its link to Similarity Matrix Regression (SMR), followed by its extension to incorporate multiple studies.

### 2.1 Microbiome regression-based kernel association test for a single study and its connection with similarity matrix regression

When analyzing data from a single study, one popular approach is the microbiome regression-based kernel association test (MiRKAT) [19]. In this method, data are summarized as independent observations of the form (*Y*_*i*_, ***X***_*i*_, ***M***_*i*_) for *i* = 1, · · ·, *n*. Here, *Y*_*i*_ represents the outcome of interest, which can be either continuous or binary. The covariates we want to adjust for, such as age, gender, and MSM status, are included in ***X***_*i*_ = (1, *x*_*i*1_, · · ·, *x*_*ip*_)^*T*^, where *p* is the number of covariates. The abundances of all taxa/OTUs in the microbial community are represented by ***M***_*i*_ = (*m*_*i*1_, · · ·, *m*_*iq*_), where *q* is the total number of taxa/OTUs.

MiRKAT compares pairwise similarity in the outcome variable to pairwise similarity in the microbiome profiles, with a high correspondence suggesting an association. It employs a kernel machine regression framework [22, 23, 24] to relate covariates and microbiome profiles to outcomes. Specifically, for a continuous outcome, we assume a linear kernel machine regression model,

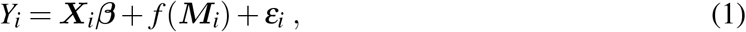

and for a binary outcome, we use the logistic kernel machine regression model,

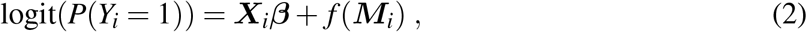

where *ε*_*i*_ is the random error with mean 0 and a common variance, *f* (·) captures the association between microbial community and the outcome, and additional covariates ***X***_*i*_ are adjusted for through a linear function. Under the null hypothesis of no microbiome effect, *f* (***M***) = 0.

In MiRKAT, *f* (***M***_***i***_) is assumed to belong to a reproducing kernel Hilbert space generated by a positive definite kernel function *K*(·,·), such that 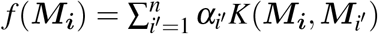. The (*i, i′*)-th element of the kernel matrix ***K***, defined as *K*(***M***_***i***_, ***M***_*i*_*′*), measures the pairwise similarity between the microbiome profiles of the *i*-th and the *i*^*′*^-th samples. To utilize the phylogenetic/taxonomic information contained in the microbiome sequences, the kernel similarity matrix can be constructed by exploiting its correspondence with the well-defined beta-diversities, which measure dissimilarities between subjects through: 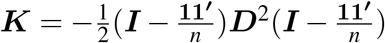, where ***D*** represents pairwise dissimilarities/distances between all subjects, calculated using beta-diversity metrics such as the Bray-Curtis or UniFrac distance. A variance component score test is then calculated through the connection between the kernel machine regression model and linear mixed models [19]: 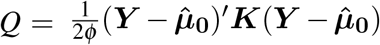, where 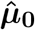 are the fitted values of ***Y*** under the null, and *φ* is the dispersion parameter.

An alternative approach to investigate the association between the outcome variable and a set of “-omics” markers is similarity matrix regression (SMR) [25]. In microbiome association studies, the SMR model can be written as

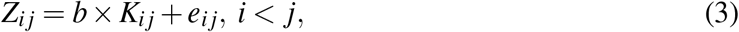

where *Z*_*i j*_, the trait similarity between subject *i* and *j*, is defined as the outer product of trait residuals, 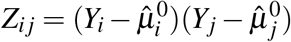, and 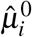 is the fitted values of *Y*_*i*_ under the null hypothesis. *K*_*i j*_ is the *i, j*-th element of microbial kernel similarity matrix. Intuitively, if the outcome is associated with microbial profiles, subjects with higher outcome similarity should also have higher microbiome similarity.

Under the null, assuming that all *Y*_*i*_’s are independent, the score statistic for testing in SMR is 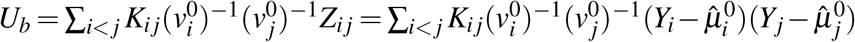, where 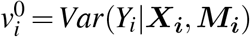 under the null. This statistic can be written in quadratic form as: 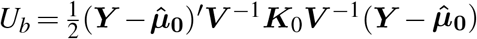, where ***K***_0_ is a matrix with diagonal elements equal to 0 and (*i, j*)-th off-diagonal elements equal to *K*_*ij*_. Under the null, *U*_*b*_ approximately follows a weighted chi-squared distribution, which can be used to assess the statistical significance.

Formulating SMR in vector form, we have

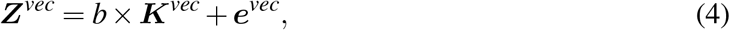

where ***Z***^*vec*^ and ***K***^*vec*^ are stacked vectors of *Z*_*i j*_, *K*_*i j*_ with all available pairwise distances (*i, j*). There is a close connection between MiRKAT and SMR [26]. Specifically, the square of the empirical sample correlation between outcome similarity and microbiome similarity in SMR, denoted by *R*^2^, is proportional to the score statistic in MiRKAT. Note that the *R*^2^ statistic represents the proportion of variation explained in SMR.

Although SMR is statistically equivalent to MiRKAT, the vector form of SMR allows for integration of multiple studies when between-study similarity matrices are not available. This is because we can exclude pairs with missing distances from the similarity vectors ***Z***^*vec*^ and ***K***^*vec*^.

### 2.2 SMRmix for integrating multiple microbiome datasets

Suppose we have a total of *K* microbiome studies. The data for the *i*-th subject in the *k*-th study is denoted as (*Y*_*ik*_, ***X***_*ik*_, ***M***_*ik*_), where *i* = 1, · · ·, *n*_*k*_, *k* = 1, · · ·, *K*. Here, *n*_*k*_ represents the number of subjects in *k*−th study, and 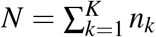 denotes the total number of subjects across *K* studies. We assume all studies are independent and do not share any subjects, and that all samples are independent within each study. A common violation of the second assumption occurs when multiple samples are longitudinal measurements from the same participant or are clustered within families in a single study. Thus, we assume that samples are clustered by studies, without additional clustering structures, although our method can be extended to allow for within-study correlations.

When microbiome data from different studies are processed through distinct protocols, subjects from these studies are not directly comparable, preventing calculation of between-study beta diversities. As a result, only intra-study microbial dissimilarities are available for association testing. Suppose the similarity matrix for all subjects across all studies (assuming it exists) is represented as follows:

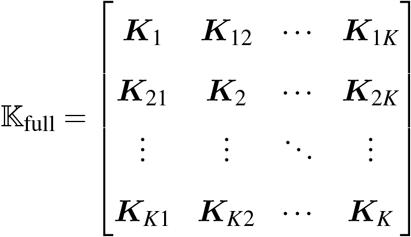

Here, the diagonal blocks ***K***_*k*_ represent the kernel similarity matrices for samples within the same study *k*, and the off-diagonals represent the microbiome similarity for pairs of samples from different studies. For existing kernel-based/distance-based methods, all elements in 𝕂_full_ must be calculated using the same distance; otherwise, the absolute values of pairwise kernels using different metrics are not on the same scale and thus not comparable.

In practice, we often only have access to the diagonal blocks, represented as follows:

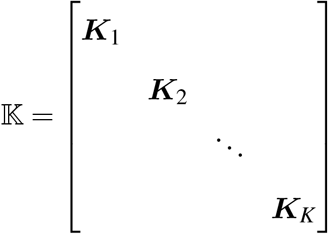

In this matrix, only diagonal blocks of 𝕂 are obtained, while the cells outside the diagonal blocks are missing. Having only the elements in diagonal blocks gives us the flexibility to choose different kernels for each study. For example, ***K***_1_ may be a UniFrac kernel while ***K***_2_ is a Bray-Curtis kernel. While it is advantageous to interpret the results when all kernels involved are of the same type, using different kernels for each study remains valid. This flexibility is particularly helpful when kernels of the same type are not available for some studies. For example, we may be able to construct a UniFrac kernel for study 1, but not for study 2 if the necessary phylogenetic tree information is unavailable, such as when using publicly available data. In this paper, we will focus on the situations where all individual ***K***_*k*_ are of the same type for simplicity. The vectorized SMR model enables us to test the association based on these blocks without the need of the off diagonal-blocks, by defining the similarity regression model only for existing *K*_*i j*_ values.

To motivate the SMR model when integrating multiple microbiome datasets, we first evaluate its counterpart model within the kernel machine regression framework, assuming the overall microbiome kernel similarity matrix 𝕂_full_ is available. For continuous outcomes, the correlated sequence kernel association test (CSKAT) [20], an extension of MiRKAT, can be used to test the association between microbiome profiles and outcomes when data from the same study are clustered. CSKAT assumes a linear mixed-effects model with study-specific random effects to link the outcome and microbial composition:

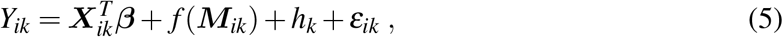

where *h*_*k*_ represents the study-specific random effect, which captures within-study correlation, and *h*_*k*_ ∼ *N*(0, *σ*^2^). The subject-level random effect is 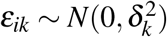.

For dichotomous outcomes, another extension of MiRKAT, GLMM-MiRKAT [21], incorporates a study-specific random effect:

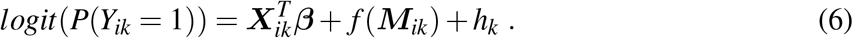

Putting samples from all studies together in vector format, we have ***h*** ∼ *N*(0, *σ* ^2^**Σ**_***h***_),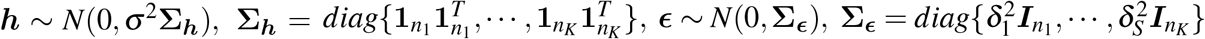.

If a full kernel matrix across all studies is available, the kernel machine regression model in CSKAT and GLMM-MiRKAT is equivalent to the (generalized) linear mixed model with ***f*** ∼ *N*(0, *τ*𝕂_full_). Hypothesis testing for *H*_0_ : *τ* = 0 can be conducted as described in the literature. However, because only the diagonal blocks ***K***_*k*_ of the overall kernel matrix are available, CSKAT and GLMM-MiRKAT cannot be applied. However, we can still calculate the trait similarity ***Z***_*k*_ and the variance-covariance matrix ***V***_*k*_ = Var(***Z***_*k*_|***X***_*k*_, ***M***_*k*_) (please refer to section A of the Supplementary Materials).

When only 𝕂 is available, we propose the following mixed effect similarity matrix regression model (SMRmix):

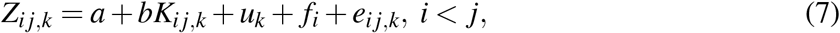

where, for subjects *i, j* in study *k*, the outcome similarity *Z*_*i j,k*_ is linked to microbiome similarity *K*_*i j,k*_. Here, *e*_*i j,k*_ is a random error that is allowed to differ across studies, *u*_*k*_ is a study-specific random effect, *f*_*i*_ is a random effect that captures the additional correlation between two pairs of trait similarities if the two pairs share a common sample. For example, *f*_*i*_ captures the additional correlation between *Z*_*i j,k*_ and *Z*_*i j*_*′,k* compared to *Z*_*i j,k*_ and *Z*_*i*_*′ j′,k*, with *i* ≠ *i*^*′*^ and *j* ≠ *j*^*′*^. Compared to the original SMR model, which is equivalent to a kernel machine regression model, our model is based on a mixed effect kernel machine regression model, thus the name SMRmix. All random effects have mean 0 and constant variance. The mathematical relationship between the random effects in SMRmix and the random effects in CSKAT and GLMM-MiRKAT is specified in Section B of Supplementary Materials.

To test whether there is a microbiome effect, we test *H*_0_ : *b* = 0. We propose the following score statistic:

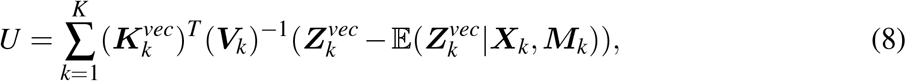

where 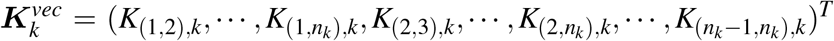 is a vector obtained by stacking all the available within-study pairwise kernels of the *k*-th study. Similarly, 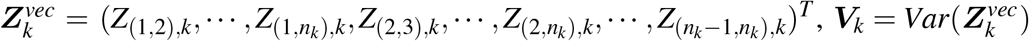 represents the corresponding outcome similarities and variances calculated under the null.

This test aggregates the score statistics from different studies, requiring only microbial similarities within studies, whereas CSKAT requires microbial pairwise similarities of all samples.

### 2.3 Perturbation approach for obtaining p-value

In the original SMR approach, the test statistic can be expressed in a quadratic form and approximated by a weighted chi-square distribution. However, for SMRmix, due to the complexity of the variance matrix 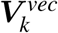 and the vectorization involved, the asymptotic distribution of *U* cannot be easily approximated. Instead, we propose a perturbation approach for obtaining the null distribution of *U* and thereby calculating the p-value. We note that, each element in 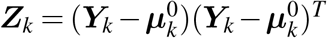 is obtained from the outer product of 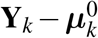, which, after some transformations, asymptotically follows a multivariate normal distribution with mean 0 and a covariance matrix that can be estimated under the null hypothesis. The idea behind the perturbation approach is to generate a large number of 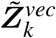 values that asymptotically have the same distribution as 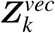. By doing so, we generate a large number of *U*^∗^ that have the same distribution as *U* under the null. Therefore, we can compare the observed test statistic with *U*^∗^ to calculate the p-value.

Perturbation procedures, sometimes referred to as resampling or Monte Carlo approaches, have been used by many researchers in various contexts [27, 28, 29, 30]. Our approach is related to these methods, but differs substantially in that we need to account for the additional correlations that arise from the clustered nature of our data. Specifically, let the estimated covariance matrix of 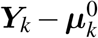 from the mixed-effect model be **Σ**_*k*_. For continuous outcomes, **Σ**_*k*_ is a symmetric matrix with diagonal elements equal to 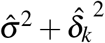 and off-diagonal elements equal to 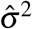. For binary outcomes, *i*−th diagonal elements is 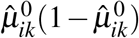, and element (*i, j*), *i*≠ *j*, in **Σ**_*k*_ is 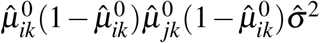.

Let 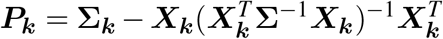, then we get 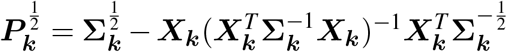.

It can be shown that, under the null hypothesis,

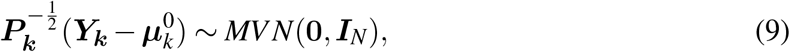

where ***I***_*N*_ is the identity matrix of size *N*.

To generate the null distribution of the test statistic, we first generate random samples 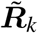 from a *k*-dimensional normal distribution with mean **0** and identity covariance matrix (i.e., the distribution of 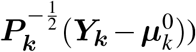. Since 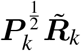 has the same distribution as 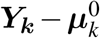, we can replace 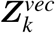 in equation (8) with a vectorized version of the outer product of 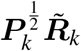, denoted as 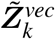. This process is repeated multiple times to construct the null distribution of the test statistic. The resulting distribution is then used to conduct hypothesis testing and calculate p-values.

### 2.4 Omnibus test

Previous research has shown that the power of statistical methods depends on whether the selected kernel effectively captures the true association signal [19]. For example, the weighted UniFrac and Bray-Curtis kernels utilize relative abundance information to construct pairwise similarities, giving higher weight to common taxa. Consequently, these kernels are more powerful when the true association involves common taxa. Conversely, the unweighted UniFrac and Jaccard kernels only use information on whether a taxon is present in a sample, giving higher weight to rare taxa, and are therefore more powerful when the true association involves rare taxa. In practice, the optimal kernel choice is unknown prior to analysis.

To address this uncertainty, we propose an omnibus test that extends SMRmix to simultaneously consider multiple kernels for each study. The omnibus-SMRmix is expected to be robust across various association patterns, losing little power compared to the best individual kernel while achieving substantially greater power than a poorly chosen kernel.

Let 𝕂_*l*_, *l* = (1,···, *L*) represent the diagonal block kernels, where the kernel for each study is of the same type, such as weighted UniFrac. Suppose there are a total of *L* such groups of kernels. For each 𝕂_*l*_, we conduct SMRmix and apply the perturbation approach as previously described to obtain a p-value. These p-values are then combined using the harmonic mean combination (HMP) [31]. In our simulations, the HMP combination demonstrated superior type I error control compared to other combination methods, such as the Cauchy combination [32], Simes combination [33] and the Tippett combination [34]. Note that even though we have *L* sets of SMRmix models, we only need to generate a single set of *B* multivariate normal distributions for perturbation across all sets of kernels. This drastically reduces computational time.

### 2.5 Simulation Studies

To assess the performance of SMRmix, we conducted extensive simulations focusing on both type I error and statistical power, comparing SMRmix against CSKAT and GLMM-MiRKAT. For power evaluation, we additionally compared SMRmix to MiRKAT-meta, which employs the HMP combination method to integrate p-values from individual MiRKAT analyses into a single p-value. We performed two sets of simulations.

#### 2.5.1 Simulation A: Microbiome data processed via the same pipeline with a common phylogenetic tree

In Simulation **A**, we generated microbiome data using a Dirichlet-Multinomial distribution, with hyperparameters estimated from real data. This simulation framework has been used in previous studies [19, 20, 35]. To account for differences in data collection across studies, we introduced a multiplicative and differential bias to capture inter-study heterogeneity. This approach ensured that the microbiome data from all studies shared the same tree, allowing the computation of a common kernel matrix that methods like CSKAT or GLMM-MiRKAT can be applied. However, with differential bias across studies, the kernel components across different studies were not directly comparable, potentially impacting the validity of results from CSKAT or GLMM-MiRKAT. This setup resembles scenarios where raw sequencing data are available and can be processed through a unified bioinformatic pipeline.

Panel A in Figure 1 illustrates the general workflow of this simulation setup. Specifically, we first generated a large OTU table via a Dirichlet-Multinomial distribution with hyperparameters estimated from a real throat microbiome dataset [36], which contained 60 samples with 856 OTUs. These generated microbiome data were treated as the “true” microbiome abundance, from which we simulated the outcome variable. The association effect was set to either the same or different across studies depending on the different simulation scenarios. The total number of counts for each sample was set to 10^6^, representing the total number of 16s rRNA molecules in the community. We denote this “true” OTU table as ***M***^∗^, which is unobserved. From this community, we conducted sub-sampling to generate the observed microbiome OTU table (denoted as ***M***), which was used for association testing.

**Figure 1:**
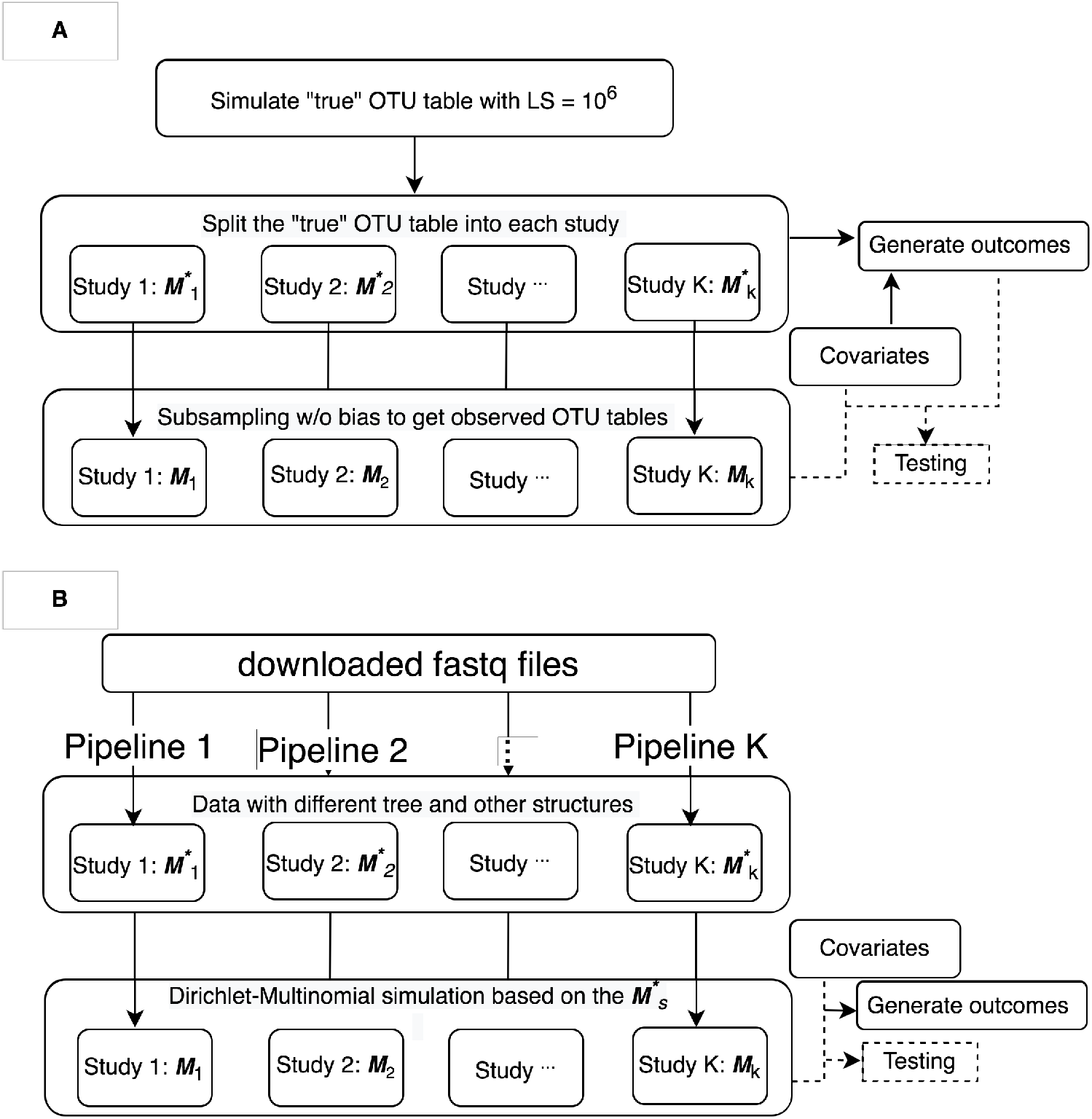
Workflow for simulation studies. Panel A: simulation scenario A, where microbiome data are processed from the same pipeline with a common tree structure. Panel B: simulation scenario B, where microbiome data are processed from different pipelines with different tree structures.

After simulating the true OTU table, we partitioned the 856 taxa into 20 clusters based on cophenetic distances calculated from the phylogenetic tree. Using the simulated microbial community, for a continuous trait, we generated the outcomes via the following model:

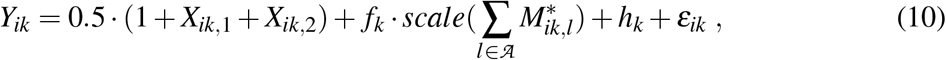

where, *i* = 1, · · ·, *n*_*k*_, *k* = 1, · · ·, *K*. The variables *X*_·,1_ and *X*_·,2_ are covariates, with *X*_·,1_ ∼ *Bernoulli*(0.5) and *X*_·,2_ ∼ *N*(0, 1). Here, 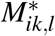 is the unobserved “true” count of *l*-th taxon for individual *i* in *k*-th study, *l* ∈ 𝒜 indicates whether taxon *l* belongs to a specific taxon cluster 𝒜. The function *scale*(·) standardizes vector 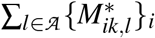 to have zero mean and unit variance. Since the total number of counts is set at 10^6^ for all samples, this is analogous to using the true relative abundances. The parameter *f*_*k*_ represents the effect size for *k*-th study. Between-study variation and within-study variation are described by *h*_*k*_ ∼ *N*(0, *σ* ^2^),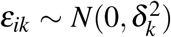, respectively. We set *σ* ^2^ = 1 and the same 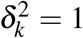 for all *k*, or different combinations of 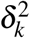 using 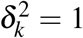, for *k* = 1, 2, 3, 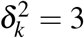, for *k* = 4, 5, 6, 7, 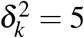, for *k* = 8, 9, 10 to evaluate type I error. We set *n*_*k*_ = 50, *K* = 10 so that we have a total of 10 studies with each study having 50 study subjects.

For binary outcome, we generated outcomes by

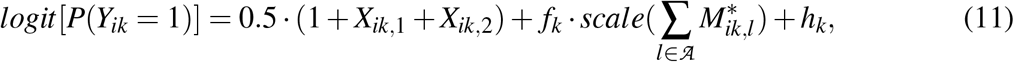

in which *X*_·,1_, *X*_·,2_, 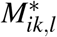 and *h*_*k*_ are defined the same way as before.

For type I error evaluation, we simulated data with *f*_*k*_ = 0 for all *k*, indicating no taxon was associated with the outcome in any study. For power evaluation, we considered two scenarios:

**Scenario A1** the microbiome effect is the same across all studies.

**Scenario A2** the microbiome effects differs across studies.

For scenario A1, we set *f*_*k*_ = *c* for all *k* and 𝒜 was the 16th cluster, which was a common cluster representing 19.6% of the total true OTU reads. The constant *c* was varied to assess the power. In scenario A2, we set *f*_*k*_ = *c* for *k* = 1, · · ·, 5 and *f*_*k*_ = −*c* for *k* = 6, · · ·, 10. Other clusters had no association with the outcome *Y*.

This outcome generation process linked the response variable to the “true” microbiome community, which is unobserved in real settings. We then generated the observed OTU table (***M***) by sub-sampling from the “true” OTU table, simulating the sequencing process as a random sampling (with or without bias) from the 16S rRNA molecules (a total of 10^6^ per sample). Two methods of random sub-sampling were considered. The first method is unbiased sampling. In this method, each molecule had an equal probability of being sampled, representing an unbiased sequencing process. We fixed the library size for each sample at 1,000 reads and generated observed OTU counts from a multinomial distribution with probabilities 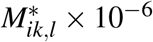 and total read counts of 1,000. In this case, the observed OTU counts were a faithful representation of the underlying community with only random sampling errors. The second method is biased sampling, where we introduced bias into the sequencing process. Some taxa were observed with higher or lower relative abundances compared to the true community. First, we generated the observed OTU table as in the unbiased sampling method. Then, for the first five studies, we inflated the read counts of the most abundant taxa by 1.1-fold. These “common” taxa were defined as the top 10% with the highest relative abundances in the true community. Consequently, the common taxa were over-represented, while other taxa had lower relative abundances, introducing bias into the observed data. Due to this bias, the beta-diversities between samples from different studies were not reliable.

All hypothesis testings and model evaluations were conducted using these “observed” datasets, as they reflect real-world data scenarios. We applied SMRmix (including both single-kernel and omnibus versions), CKSAT (for continuous outcomes), and GLMM-MiRKAT (for binary outcomes). MiRKAT-meta was also included in power evaluation. Each evaluation involved 10,000 replicates for type I error assessment and 5,000 replicates for power evaluation. We tested the null hypothesis of no association between the microbiome profiles and outcomes. For the tests with a single kernel, we used three kernels: weighted UniFrac(*K*_*w*_), unweighted UniFrac(*K*_*u*_) and Bray-Curtis (*K*_*bc*_). The UniFrac kernels utilize phylogenetic information while the Bray-Curtis does not. For SMRmix-Omnibus test, we synthesized test results from five kernelsm including weighted UniFrac, unweighted UniFrac, Bray-Curtis, and generalized UniFrac with weight parameter of 0 and 0.5.

#### 2.5.2 Simulation B: Microbiome data processed through different pipelines with different tree structures

In simulations (**B**), we reprocessed the raw sequencing (FASTQ) files from a real microbiome study through different bioinformatic pipelines, resulting in multiple OTU tables with distinct tree structures. For each dataset processed through a distinct bioinformatic pipelines, we estimated hyperparameters of the Dirichlet-Multinomial distribution and subsequently simulated additional microbiome data with its corresponding tree structure. This set of simulations reflects scenarios where the collected microbiome datasets are pre-processed through different bioinformatic pipelines, but access to the raw sequencing data is unavailable, a common situation when working with public datasets. In such scenarios, generating a single kernel matrix is not feasible, and approaches that require a common kernel matrix, such as CSKAT and GLMM-MiRKAT, cannot be applied.

Panel B in Figure 1 summarizes the general workflow of this simulation. The raw sequencing files were from a real gut microbiome study [10], which contains stool microbiome data from 82 HIV-positive and 40 HIV-negative individuals. The samples underwent PCR amplification of the V4 region and were sequenced on the Illumina MiSeq platform. We downloaded the paired-end sequencing data in FASTQ format from Synapse (ID: syn22273201), and reprocess it through different pipelines to generate a total of nine dataset, each with distinct phylogenetic tree structures and OTU definitions.

Among the nine datasets, six of them were processed through the QIIME2 pipeline [37]. Within QIIME2, paired-end sequencing data were joined and filtered, and two denoising techniques-DADA2 [38] and Deblur [39]-were used along with three phylogeny construction approaches-FastTree [40], RAxML [41] and IQ-TREE [42]. This combination resulted in six (2 *×* 3) distinct datasets. The remaining three datasets were generated using the original QIIME pipeline [43], employing three different OTU-picking strategies: closed-reference OTU picking using SILVA (Release 132) [44, 45], closed-reference OTU picking using Greengenes (Version 13 8) [46], and De novo OTU picking. Due to differences in the phylogenetic trees and taxa definitions across these processing steps, a common kernel matrix was not available in this simulation. This represents a scenario in which multiple public datasets are integrated without access to the raw sequencing data.

After generating the nine datasets, we partitioned the taxa in each dataset into 20 clusters. We then simulated new count tables based on these datasets using Dirichlet-Multinomial distribution and the hyperparameters estimated from the nine count tables. The outcomes were generated using equation (10) for continuous traits and equation (11) for binary traits. Similar to scenario A, we evaluate type I error and power using *σ* ^2^ = 1 and either the same 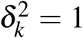 for all *k*, or different combinations of 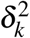 where 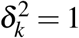 for *k* = 1, 2, 3, 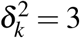 for *k* = 4, 5, 6, 7, 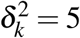 for *k* = 8, 9, 10. To assess power, we assumed that the microbiome effect was consistent across the nine tables, because they originated from the same microbiome community. We specified *f*_*k*_ = *c* for *k* ∈ *{*1,···, 9*}* and selected the 16th cluster in each dataset as 𝒜.

Unlike Simulation A, we did not introduce a process to simulate bias in this scenario, as there was no “true” community available for comparison. We applied the proposed SMRmix and metaanalysis procedures to the simulated data, noting that CSKAT and GLMM-MiRKAT could not be used due to the lack of a common kernel matrix.

## 3 Results

### 3.1 Simulation results

Table 1 summarizes the type I error rates for continuous and binary outcomes across various scenarios in simulation setups A and B. SMRmix controls type I error rates effectively across all scenarios. In simulation setup A, CSKAT and GLMM-MiRKAT also showed valid type I error rates, regardless of sequencing bias. For binary outcomes, SMRmix was slightly conservative.

**Table 1:**
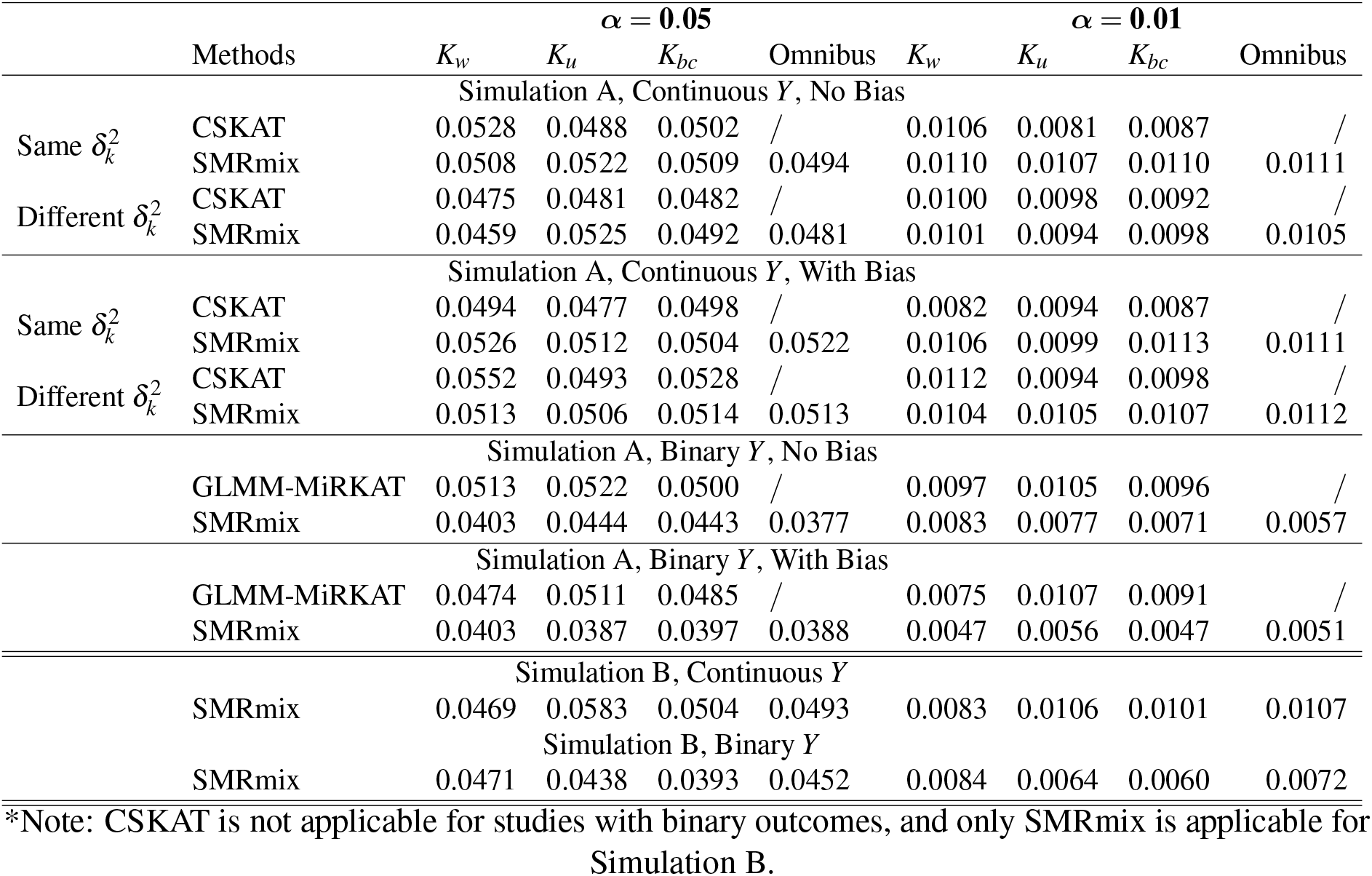
Empirical type I error rates comparing CSKAT, GLMM-MiRKAT, and SMRmix at different *α* cutoffs (0.05 and 0.01) in simulations. For continuous outcome with the same 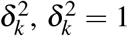,*∀k*; for continuous outcome with different 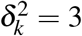 for *k* = 1, 2, 3, *δ*^2^ = 3 for *k* = 4,6,5, 7, 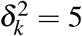 for *k* = 8, 9, 10. For binary scenarios, *σ* ^2^ = 1.

Figure 2 presents the power results for simulation setups A and B without sequencing bias. In simulation A1 with both continuous and binary outcomes, (left panels in Figure 2) when all studies share the same microbiome effect and there is no sequencing bias, CSKAT/GLMM-MiRKAT achieves higher power than SMRmix when comparing methods using the same type of kernel (e.g., Bray-Curtis kernel). This is expected, as in this scenario, microbiome data across studies are comparable and CSKAT/GLMM-MiRKAT leverage the similarities between samples from different studies, offering more efficient use of the data. In contrast, the meta-analysis approach that combines p-values from multiple MiRKAT models exhibits substantial power loss.

**Figure 2:**
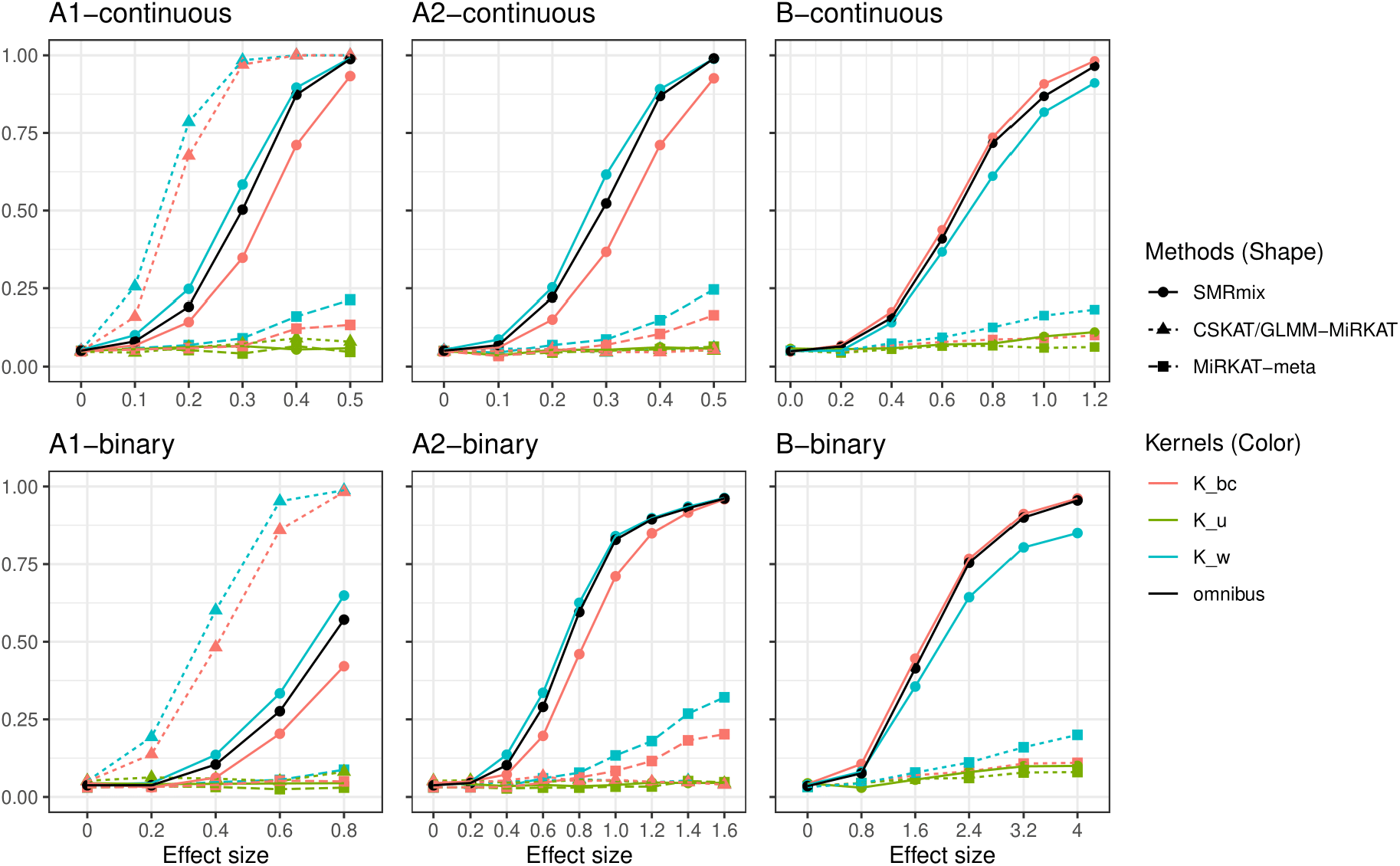
Statistical power for simulations A and B when there is no bias in sequencing sampling. For continuous scenarios (upper three panels), 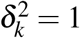, *∀k*, and *σ* 2 = 1.

When comparing different kernels within the same method, both CSKAT and SMRmix demonstrate that the weighted UniFrac is the most powerful, followed by Bray-Curtis, and the unweighted UniFrac is the least powerful. This is because the microbiome effect in this scenario targets a phylogenetically-informed cluster of common taxa. A similar trend is observed even when sequencing bias is introduced (left panels in Figure S1 in Supplementary Materials), likely due to the limited magnitude of bias introduced in this simulation.

In scenarios A2, where studies have distinct microbiome effects (middle panels in Figure 2) and no sequencing bias, CSKAT/GLMM-MiRKAT, regardless of the kernel choice, show minimal power. This is because CSKAT/GLMM-MiRKAT implicitly assume that the microbiome-outcome relationship is consistent across studies via the same kernel function, which is violated in this scenario. In contrast, SMRmix demonstrates higher power. While the power of meta-analysis is substantially lower than SMRmix, it still performs better than CSKAT/GLMM-MiRKAT in this scenario. When comparing different kernels, the results are similar to those observed in Scenario A1. Notably, SMRmix maintains its power effectively between Simulations A1 and A2, demonstrating its robustness in scenarios with differing effect sizes across studies. A similar conclusion is drawn when sequencing bias is introduced (right panels of Figure S1 in the Supplementary Materials).

In simulation setup B, SMRmix exhibits much higher power than the meta-analysis procedure (right panels in Figure 2). CSKAT and GLMM-MiRKAT are not applicable in this simulation.

The power results for continuous outcomes with varying variances of study-specific random effects 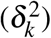 are displayed in Figure S2 of the Supplementary Materials. The conclusions of performance under scenarios A1 and A2 remain consistent with previous findings. Notably, in simulation scenario A2, the meta analysis suffers a substantial power loss, while SMRmix stands out as the only method capable of effectively detecting associations.

### 3.2 Application to HIV studies

We utilized SMRmix to analyze data from the HIV Microbiome Re-analysis Consortium [47], which re-analyzed *α*−diversity using raw 16S rRNA sequencing data from 17 studies. These studies aimed to characterize the gut dysbiosis in HIV+ individuals. The microbiome amplicon sequencing data and metadata of these studies are available on Synapse (ID: syn18406854) and NCBI. The 17 studies differ in their study design, institution of participants recruitment, study population, type of samples and various factors in the sequencing protocol including the 16s rRNA variable regions that were sequenced and the sequencing platform. The consortium reprocessed the raw sequencing data using standardized protocols and conducted taxonomic assignments in species level through Resphera Insight.

Our goal was to evaluate the association between microbiome composition and HIV status, MSM status by integrating all these studies and draw a consistent conclusion. Out of the 17 datasets, 14 studies(763 samples and 666 taxa) have subject-level information on MSM status, age, and HIV status, which we used for further analysis. Because a common phylogenetic tree is not available from Synapse webpage, we created a phylogentic tree using the “tax tree” function in R package “LTN” based on the taxonmy relationship [48]. The Principal Coordinates Analysis (PCoA) plots, which visualize the distance matrix between study subjects, are presented in Supplementary Figure S3.

We conducted the association tests utilizing single-kernel SMRmix based on kernels including weighted UniFrac(*K*_*w*_), unweighted UniFrac(*K*_*u*_), Jaccard(*K*_*jac*_), Bray-Curtis(*K*_*bc*_) and omnibus SMRmix using the above kernels. Suggested by previous studies [3], we adjusted HIV status when the outcome variable is MSM, and adjusted MSM when the outcome variable is HIV status.

The statistical model was established as

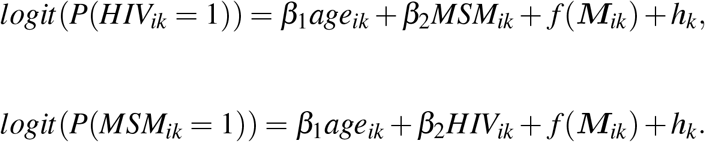

We also conducted an analysis in which each individual sample was analyzed separately, and HMP combination was used to obtain the final p-values, similar to the approach used in the simulation studies. The p-values are reported in Table 2. MiRKAT-meta yielded a p-value of 0.0623 for the outcome of HIV, while SMRmix reported a p-value *<* 0.0001, in line with our simulation results.

**Table 2:** P-values of using SMRmix to test associations between microbiome composition and outcome variables HIV Status and MSM adjusting for age.

|  | Outcome | SMRmix |  |  |  |  | MiRKAT-meta |
| --- | --- | --- | --- | --- | --- | --- | --- |
| | | $K_w$ | $K_u$ | $K_{jac}$ | $K_{bc}$ | Omnibus | Omnibus |
| HIV data | HIV | $< 0.0001$ | $< 0.0001$ | $< 0.0001$ | $< 0.0001$ | $< 0.0001$ | 0.0623 |
| | MSM | $< 0.0001$ | $< 0.0001$ | $< 0.0001$ | $< 0.0001$ | $< 0.0001$ | $< 0.0001$ |
| CRC data | CRC | $< 0.0001$ | $< 0.0001$ | $< 0.0001$ | $< 0.0001$ | $< 0.0001$ | $< 0.0001$ |

For the outcome HIV status, SMRmix with weighted UniFrac, UniFrac, Jaccard, and Bray-Curtis kernels all discovered significant signals (p-values *<* 0.0001). Synthesizing the results based on above kernels, omnibus SMRmix found HIV status and microbiome composition associated (p-value *<* 0.0001). For the outcome MSM, similar conclusions were obtained that SMRmix with all kernels show significant connection between MSM and microbiome composition.

### 3.3 Application to colorectal cancer data

We applied the proposed method to eleven metagenomic datasets containing species-level OTU data for fecal microbiomes of individuals, consisting of 354 colorectal cancer patients and 393 healthy controls. These datasets were obtained from the curatedMetagenomicData package in Bioconductor, which provides standardized microbial profiling using MetaPhlAn [49]. These studies exhibit great heterogeneity in sequencing platforms (Illumina HiSeq, Illumina NextSeq), DNA extraction kits (Gnome, MoBio, Qiagen), and other technical factors. The sample sizes varied from 9 to 279 across the studies (Supplementary Table S1). PCoA plots are provided in Supplementary Figure S4. Samples from the same study tend to cluster together, with noticeable shifts across clusters, suggesting study-specific variation.

The objective was to evaluate differences in microbiome composition between colorectal cancer patients and healthy controls. Patient demographics are summarized in Supplementary Table S1. To examine variations in patient demographics within studies, we fit a generalized linear mixed model (GLMM), using disease status as the outcome and age, gender, and BMI as control variables. Significant variations were found in patient demographics within studies, including age (GLMM p-value *<* 0.0001), gender (GLMM p-value = 0.0087), and BMI (GLMM p-value = 0.0297). To account for these variations, we adjusted for these factors when applying SMRmix to integrate information from these studies. Using SMRmix, we found a significant difference in microbiome composition between colorectal cancer patients and healthy controls based on weighted UniFrac, unweighted UniFrac, Jaccard, and Bray-Curtis kernels (See Table 2), and MiRKAT-meta yielded similar results. These findings are consistent with previous research, which also suggests a distinct microbiome profile associated with colorectal cancer [50].

## 4 Discussion

In this paper, we present a novel approach, SMRmix for testing the association between microbiome composition and outcomes of interest, by combining the concepts of similarity matrix regression and kernel machine regression. This method is particularly beneficial in the context of integrative analysis of microbiome data from multiple studies, cohorts or protocols, where interstudy pairwise distances are either absent or unreliable. SMRmix effectively addresses the limitations of existing kernel regression tests by leveraging only within-study pairwise distances for testing. In simulations, SMRmix showed well-controlled Type I error rates. We compared SMRmix to two other methods, CSKAT and GLMM-MiRKAT (which require the usually non-available between-study pairwise distance), for microbiome compositional analysis. Our results showed that if distances of all pairs of samples are available and when the microbiome effect is homogeneous, SMRmix has relatively lower power. However, SMRmix exhibits superior performance in scenarios where the effects of taxa are a mixture of positive and negative effects, and when different effect sizes result in a significant reduction in power for CSKAT and GLMM-MiRKAT. This highlights the robustness and effectiveness of SMRmix, particularly when inter-study pairwise distances may not be reliable or available. We also compared SMRmix with a meta-analysis procedure (which, like SMRmix, doesn’t require the between-study pairwise distance), and showed higher power of SMRmix across all simulation setups. In summary, our proposed SMRmix method represents a novel and effective approach for microbiome compositional analysis, with the potential to contribute to ongoing efforts to better understand the complex interactions between microbial communities and host health.

Kernel selection is a critical challenge for any kernel-based approach, as choosing an unsuitable kernel can severely reduce the statistical power despite controlling the type I error. In microbiome studies, each kernel function assumes a distinct microbiome effect in the community, and improper choice of kernel can compromise the analysis [51]. To address this issue, SMRmix provides an omnibus test that integrates multiple kernel matrices to enhance the robustness of the analysis. SMRmix also allows for the use of different kernels for each study, such as Bray-Curtis distances in one study and UniFrac distances in another, thereby offering greater flexibility depending on the data availability. Our simulations demonstrate that single-kernel tests can be sensitive to kernel selection. In contrast, our omnibus SMRmix approach synthesizes results from multiple kernel-based tests by combining p-values and only incurs a small loss of power relative to using the best kernel. By contrast, the omnibus test improves the power substantially compared to a poor kernel choice. Given that the best kernel is often unknown in practice, the omnibus test is a practical and useful alternative.

In applying SMRmix to investigate the relationship between human gut microbiome composition and HIV infection using data from 14 studies, we aimed to address conflicting findings in the literature. Although many studies have reported gut microbiome dysbiosis associated with HIV infection and highlighted the potential role of “leaky gut” in HIV pathogenesis [1, 2, 3, 4, 5, 6, 7, 8, 9, 10, 11], others have suggested that MSM status may confound the HIV-microbiome relationship [3]. Our analysis adjusted for age and MSM status to assess the association between microbiome composition and HIV status, and similarly adjusted for age and HIV infection to examine the association between microbiome composition and MSM status. By integrating information from multiple studies on this topic, we found significant associations between microbiome composition and HIV (adjusting for age and MSM) and between microbiome and MSM (adjusting for age and HIV). These findings provide compelling evidence of HIV-related dysbiosis that is independent of MSM status, challenging the notion that MSM is only a confounding factor in the HIV-microbiome relationship.

## Supplementary Information

### Supplementary Files

Supplemental files can be obtained through online version of the paper at *Microbiome*.

## Authors’ contributions

MH contributed to the development of the method, performed simulation studies and comparisons, and wrote the manuscript. NZ conceived the study, primarily developed the method, and wrote the manuscript. All authors read and approved the final manuscript.

## Availability of data and materials

The implementation of SMRmix in R codes are available in the GitHub repository https://github.com/mengyu-he/SMRmix.

## Ethics approval and consent to participate

Not applicable.

## Consent for publication

Not applicable.

## Acknowledgements

Not applicable.

## Author affiliations

Department of Biostatistics and Bioinformatics, Emory University, Atlanta, 30322, GA, USA (Mengyu He)

Department of Biostatistics, Johns Hopkins University, Baltimore, MD 21205, USA (Ni Zhao)

## References

[1] Dubourg, G. et al. Gut microbiota associated with HIV infection is significantly enriched in bacteria tolerant to oxygen. BMJ Open Gastroenterology 3, e000080 (2016). URL https://bmjopengastro.bmj.com/lookup/doi/10.1136/bmjgast-2016-000080.

[2] Mutlu, E. A. et al. A Compositional Look at the Human Gastrointestinal Microbiome and Immune Activation Parameters in HIV Infected Subjects. PLoS Pathogens 10, e1003829 (2014). URL https://dx.plos.org/10.1371/journal.ppat.1003829.

[3] Noguera-Julian, M. et al. Gut Microbiota Linked to Sexual Preference and HIV Infection. EBioMedicine 5, 135–146 (2016). URL https://linkinghub.elsevier.com/retrieve/pii/S2352396416300287.

[4] Nowak, P. et al. Gut microbiota diversity predicts immune status in HIV-1 infection. AIDS 29, 2409–2418 (2015). URL https://journals.lww.com/00002030-201511280-00004.

[5] Pinto-Cardoso, S. et al. Fecal Bacterial Communities in treated HIV infected individuals on two antiretroviral regimens. Scientific Reports 7, 43741 (2017). URL http://www.nature.com/articles/srep43741.

[6] Villanueva-Millán, M. J., Pérez-Matute, P., Recio-Fernández, E., Lezana Rosales, J. M. & Oteo, J. A. Differential effects of antiretrovirals on microbial translocation and gut microbiota composition of HIV-infected patients: Villanueva-Millán MJ et al. Journal of the International AIDS Society 20, 21526 (2017). URL http://doi.wiley.com/10.7448/IAS.20.1.21526.

[7] Yu, G., Fadrosh, D., Ma, B., Ravel, J. & Goedert, J. J. Anal microbiota profiles in HIV-positive and HIV-negative MSM. AIDS 28, 753–760 (2014). URL https://journals.lww.com/00002030-201403130-00013.

[8] Sun, Y. et al. Fecal bacterial microbiome diversity in chronic HIV-infected patients in China. Emerging Microbes & Infections 5, 1–7 (2016). URL https://www.tandfonline.com/doi/full/10.1038/emi.2016.25.

[9] Vázquez-Castellanos, J. F. et al. Altered metabolism of gut microbiota contributes to chronic immune activation in HIV-infected individuals. Mucosal Immunology 8, 760–772 (2015). URL http://www.nature.com/articles/mi2014107.

[10] Monaco, C. et al. Altered Virome and Bacterial Microbiome in Human Immunodeficiency Virus-Associated Acquired Immunodeficiency Syndrome. Cell Host & Microbe 19, 311–322 (2016). URL https://linkinghub.elsevier.com/retrieve/pii/ S193131281630052X.

[11] Vesterbacka, J. et al. Richer gut microbiota with distinct metabolic profile in HIV infected Elite Controllers. Scientific Reports 7, 6269 (2017). URL http://www.nature.com/articles/s41598-017-06675-1.

[12] Volpe, G. E. et al. Associations of Cocaine Use and HIV Infection With the Intestinal Microbiota, Microbial Translocation, and Inflammation. Journal of Studies on Alcohol and Drugs 75, 347–357 (2014). URL http://www.jsad.com/doi/10.15288/jsad.2014.75.347.

[13] Ling, Z. et al. Alterations in the Fecal Microbiota of Patients with HIV-1 Infection: An Observational Study in A Chinese Population. Scientific Reports 6, 30673 (2016). URL http://www.nature.com/articles/srep30673.

[14] Dinh, D. M. et al. Intestinal Microbiota, Microbial Translocation, and Systemic Inflammation in Chronic HIV Infection. Journal of Infectious Diseases 211, 19–27 (2015). URL https://academic.oup.com/jid/article-lookup/doi/10.1093/infdis/jiu409.

[15] Nowak, R. G. et al. Rectal microbiota among HIV-uninfected, untreated HIV, and treated HIV-infected in Nigeria. AIDS 31, 857–862 (2017). URL https://journals.lww.com/00002030-201703270-00014.

[16] Lozupone, C. et al. Alterations in the Gut Microbiota Associated with HIV-1 Infection. Cell Host & Microbe 14, 329–339 (2013). URL https://linkinghub.elsevier.com/retrieve/pii/S1931312813002886.

[17] McArdle, B. H. & Anderson, M. J. Fitting multivariate models to community data: a comment on distance-based redundancy analysis. Ecology 82, 290–297 (2001).

[18] MetaHIT Consortium (additional members) et al. Enterotypes of the human gut microbiome. Nature 473, 174–180 (2011). URL http://www.nature.com/articles/nature09944.

[19] Zhao, N. et al. Testing in microbiome-profiling studies with MiRKAT, the microbiome regression-based kernel association test. The American Journal of Human Genetics 96, 797–807 (2015). URL 10.1016/j.ajhg.2015.04.003.

[20] Zhan, X. et al. A small-sample kernel association test for correlated data with application to microbiome association studies. Genetic Epidemiology 42, 772–782 (2018). URL https://onlinelibrary.wiley.com/doi/abs/10.1002/gepi.22160.

[21] Koh, H., Li, Y., Zhan, X., Chen, J. & Zhao, N. A Distance-Based Kernel Association Test Based on the Generalized Linear Mixed Model for Correlated Microbiome Studies. Frontiers in Genetics 10, 458 (2019). URL https://www.frontiersin.org/article/10.3389/fgene.2019.00458/full.

[22] Kwee, L. C., Liu, D., Lin, X., Ghosh, D. & Epstein, M. P. A Powerful and Flexible Multilocus Association Test for Quantitative Traits. The American Journal of Human Genetics 82, 386–397 (2008). URL https://linkinghub.elsevier.com/retrieve/pii/S0002929708000888.

[23] Wu, M. C. et al. Powerful SNP-Set Analysis for Case-Control Genome-wide Association Studies. The American Journal of Human Genetics 86, 929–942 (2010). URL https://linkinghub.elsevier.com/retrieve/pii/S000292971000248X.

[24] Wu, M. et al. Rare-Variant Association Testing for Sequencing Data with the Sequence Kernel Association Test. The American Journal of Human Genetics 89, 82–93 (2011). URL https://linkinghub.elsevier.com/retrieve/pii/S0002929711002229.

[25] Tzeng, J.-Y., Zhang, D., Chang, S.-M., Thomas, D. C. & Davidian, M. Gene-Trait Similarity Regression for Multimarker-Based Association Analysis. Biometrics 65, 822–832 (2009). URL http://doi.wiley.com/10.1111/j.1541-0420.2008.01176.x.

[26] Zhan, X. Relationship Between MiRKAT and Coefficient of Determination in Similarity Matrix Regression. Processes 7, 79 (2019). URL http://www.mdpi.com/2227-9717/7/2/79.

[27] Lin, D. Y. An efficient Monte Carlo approach to assessing statistical significance in genomic studies. Bioinformatics 21, 781–787 (2005). URL https://academic.oup.com/bioinformatics/article-lookup/doi/10.1093/bioinformatics/bti053.

[28] Conneely, K. N. & Boehnke, M. So Many Correlated Tests, So Little Time! Rapid Adjustment of P Values for Multiple Correlated Tests. The American Journal of Human Genetics 81, 1158–1168 (2007). URL https://linkinghub.elsevier.com/retrieve/pii/ S0002929707637665.

[29] Chapman, J. & Whittaker, J. Analysis of multiple SNPs in a candidate gene or region. Genetic Epidemiology 32, 560–566 (2008). URL http://doi.wiley.com/10.1002/gepi.20330.

[30] Pan, W., Han, F. & Shen, X. Test Selection with Application to Detecting Disease Association with Multiple SNPs. Human Heredity 69, 120–130 (2010). URL https://www.karger.com/Article/FullText/264449.

[31] Wilson, D. J. The harmonic mean p -value for combining dependent tests. Proceedings of the National Academy of Sciences 116, 1195–1200 (2019). URL http://www.pnas.org/lookup/doi/10.1073/pnas.1814092116.

[32] Liu, Y. & Xie, J. Cauchy combination test: A powerful test with analytic p-value calculation under arbitrary dependency structures. Journal of the American Statistical Association 115 (2020).

[33] Simes, R. J. An improved bonferroni procedure for multiple tests of significance. Biometrika 73 (1986).

[34] Tippett, L. H. C. The methods of statistics Vol. 2d ed., re (Williams Norgate ltd, London, 1931).

[35] Chen, J. & Li, H. Kernel methods for regression analysis of microbiome compositional data, Vol. 55 (2013).

[36] Charlson, E. S. et al. Disordered Microbial Communities in the Upper Respiratory Tract of Cigarette Smokers. PLoS ONE 5, e15216 (2010). URL https://dx.plos.org/10.1371/journal.pone.0015216.

[37] Bolyen, E. et al. Reproducible, interactive, scalable and extensible microbiome data science using QIIME 2. Nature Biotechnology 37, 852–857 (2019). URL http://www.nature.com/articles/s41587-019-0209-9.

[38] Callahan, B. J. et al. Dada2: High-resolution sample inference from illumina amplicon data. Nature Methods 13 (2016).

[39] Amir, A. et al. Deblur rapidly resolves single-nucleotide community sequence patterns. mSystems 2 (2017). URL 10.1128/mSystems.00191-16.

[40] Price, M. N., Dehal, P. S. & Arkin, A. P. FastTree: Computing Large Minimum Evolution Trees with Profiles instead of a Distance Matrix. Molecular Biology and Evolution 26, 1641–1650 (2009). URL https://academic.oup.com/mbe/article-lookup/doi/10.1093/molbev/msp077.

[41] Stamatakis, A. RAxML-VI-HPC: maximum likelihood-based phylogenetic analyses with thousands of taxa and mixed models. Bioinformatics 22, 2688–2690 (2006). URL https://academic.oup.com/bioinformatics/article-lookup/doi/10.1093/bioinformatics/btl446.

[42] Nguyen, L.-T., Schmidt, H. A., von Haeseler, A. & Minh, B. Q. IQ-TREE: A Fast and Effective Stochastic Algorithm for Estimating Maximum-Likelihood Phylogenies. Molecular Biology and Evolution 32, 268–274 (2015). URL https://academic.oup.com/mbe/article-lookup/doi/10.1093/molbev/msu300.

[43] Caporaso, J. G. et al. QIIME allows analysis of high-throughput community sequencing data. Nature Methods 7, 335–336 (2010). URL http://www.nature.com/articles/nmeth.f.303.

[44] Quast, C. et al. The SILVA ribosomal RNA gene database project: improved data processing and web-based tools. Nucleic Acids Research 41, D590– D596 (2012). URL http://academic.oup.com/nar/article/41/D1/D590/1069277/The-SILVA-ribosomal-RNA-gene-database-project.

[45] Yilmaz, P. et al. The SILVA and “all-species living tree project (LTP)” taxonomic frameworks. Nucleic Acids Research 42, D643–D648 (2013). URL 10.1093/nar/gkt1209.

[46] McDonald, D. et al. An improved Greengenes taxonomy with explicit ranks for ecological and evolutionary analyses of bacteria and archaea. The ISME Journal 6, 610–618 (2012). URL http://www.nature.com/articles/ismej2011139.

[47] Tuddenham, S. A. et al. The impact of human immunodeficiency virus infection on gut microbiota α-diversity: An individual-level meta-analysis. Clinical Infectious Diseases 70 (2020).

[48] Wang, Z., Mao, J. & Ma, L. Microbiome compositional analysis with logistic-tree normal models. arXiv preprint arXiv:2106.15051 (2021).

[49] Pasolli, E. et al. Accessible, curated metagenomic data through ExperimentHub. Nat. Methods 14, 1023–1024 (2017).

[50] Ahn, J. et al. Human gut microbiome and risk for colorectal cancer. JNCI: Journal of the National Cancer Institute 105, 1907–1911 (2013). URL 10.1093/jnci/djt300.

[51] Plantinga, A. et al. Mirkat-s: a community-level test of association between the microbiota and survival times. Microbiome 5, 17 (2017).

