## Supplemental file for "A Mixed Effect Similarity Matrix Regression Model (SMRmix) for Integrating Multiple Microbiome Datasets at Community Level"

#### A Connection Between SMRmix and Mixed effect Kernel Machine Regression Models

When data are obtained from a single study, the corresponding kernel model regression model, as specified by equations (2.1) and (2.2) in the main manuscript, is given by $Y_i = \mathbf{X_i}\beta + f(\mathbf{M_i}) + \varepsilon_i$, and $\text{logit}(P(Y_i = 1)) = \mathbf{X_i}\beta + f(\mathbf{M_i})$, respectively. Let $Z_{ij} = (Y_i - \hat{\mu}_i^0)(Y_j - \hat{\mu}_j^0)$, where $\mu_i^0$ is the expectation of $Y_i$ under the null hypothesis. The original SMR model is given by $Z_{ij} = b \times K_{ij,k} + e_{ij},\ i < j$. The correspondence between SMRmix and kernel machine regression model has been established via Zhan (2019). Here, no intercept is needed because $Z_{ij} = (Y_i - \hat{\mu}_i^0)(Y_j - \hat{\mu}_j^0)$ has already adjusted for the effect of covariates including the intercept (Tzeng et al., 2009).

When data are obtained from multiple studies, extra terms should be considered in the similarity matrix model to account for possible extra correlations among similarities. Here, the counterpart mixed-effect kernel machine regression models (as in CSKAT and GLMM-MiRKAT, Equations (2.5) and (2.6) in the main manuscript) are,

$$Y_{ik} = \mathbf{X}_{ik}^T\beta + f(\mathbf{M}_{ik}) + h_k + \varepsilon_{ik}$$
$$logit(P(Y_{ik} = 1)) = \mathbf{X}_{ik}^T\beta + f(\mathbf{M}_{ik}) + h_k\ .$$

where $h_k \sim N(0, \sigma^2)$, $\varepsilon_{ik} \sim N(0, \delta_k^2)$, and $f(\mathbf{M}_{ik})$ depicts the microbiome effect on the outcome. Let $\gamma_{ik} = f(\mathbf{M}_{ik})$, $\gamma_k = (\gamma_{1k}, \cdots, \gamma_{n_k k}) \sim \text{MVN}(\mathbf{0}, \tau \mathbf{K}_k)$, where $\mathbf{K}_k$ is the kernel similarity matrix in $k$-th study. Under the null hypothesis, $\tau = 0$, i.e. $\gamma_{ik} = 0$ for all $i$ and $k$.

To understand the trait similarity between pairs of $Z_{ij,k} = (Y_{ik} - \hat{\mu}_{ik}^0)(Y_{jk} - \hat{\mu}_{jk}^0)$, and relate it to the corresponding microbiome similarity $K_{ij,k}$, the conditional expectation and the following four quantities of conditional variance-covariance matrix are required,

1. $\text{Var}(Z_{ij,k} | \mathbf{X}, \mathbf{M})$: the variance of trait similarity of subject pair $(i, j)$ within the $k$-th study.
2. $\text{Cov}(Z_{ij,k}, Z_{it,k} | \mathbf{X}, \mathbf{M})$: the covariance of trait similarities between subject pairs $(i, j)$ and $(i, t)$ in the $k$-th study, where the pairs share the same subject $i$ ($i \neq j \neq t$).
3. $\text{Cov}(Z_{ij,k}, Z_{st,k} | \mathbf{X}, \mathbf{M})$: the covariance of trait similarities between subject pairs $(i, j)$ and $(s, t)$ in the $k$-th study, where the pairs involve four different subjects ($i \neq j \neq s \neq t$).
4. $\text{Cov}(Z_{ij,k}, Z_{st,k'} | \mathbf{X}, \mathbf{M})$: the covariance of trait similarities between subject pairs $(i, j)$ in $k$-th study and $(s, t)$ in $k'$-th study ($i \neq j, s \neq t$).

As detailed in section B, we show that, for both the continuous and binary outcomes,

$$\mathbb{E}(Z_{ij,k} | \mathbf{X}, \mathbf{M}) = a + b K_{ij,k}, \tag{A.1}$$

where a and b are two constants, $K_{ij,k}$ is the $i, j$-th element of microbiome similarity matrix $\mathbf{K}_k$.

Building on the counterpart model, the conditional variance and covariances for continuous outcomes (See details in section B.1),

$$\begin{aligned}\mathrm{Var}(Z_{ij,k}|\mathbf{X},\mathbf{M}) =& \tau\sigma^2(K_{ii,k}+2K_{ij,k}+K_{jj,k})+\tau^2(K_{ii,k}K_{jj,k}+K_{ij,k}^2)\\ &+2\sigma^4+\tau\delta_k^2(K_{ii,k}+K_{jj,k})+2\delta_k^2\sigma^2+\delta_k^4 \qquad (A.2)\\ \mathrm{Cov}(Z_{ij,k},Z_{it,k}|\mathbf{X},\mathbf{M}) =& \tau\sigma^2(K_{it,k}+K_{ii,k}+K_{ji,k}+K_{jt,k})+\tau^2(K_{ii,k}K_{jt,k}+K_{it,k}K_{ij,k})\\ &+2\sigma^4+\tau\delta_k^2K_{jt,k}+\delta_k^2\sigma^2 \qquad (A.3)\\ \mathrm{Cov}(Z_{ij,k},Z_{st,k}|\mathbf{X},\mathbf{M}) =& \tau\sigma^2(K_{it,k}+K_{is,k}+K_{js,k}+K_{jt,k})+\tau^2(K_{is,k}K_{jt,k}+K_{it,k}K_{js,k})\\ &+2\sigma^4 \qquad (A.4)\\ \mathrm{Cov}(Z_{ij,k},Z_{st,k'}|\mathbf{X},\mathbf{M}) =& \tau^2(K_{is,kk'}K_{jt,kk'}+K_{it,kk'}K_{js,kk'}) \qquad (A.5)\end{aligned}$$

where $\delta_k^2$ denotes the variance of $\varepsilon_{ik}$, $\sigma^2$ denotes the variance of $h_k$, $K_{ij,k}$ represents that microbiome similarity between subject $i$ and subject $j$ within the study $k$, and $K_{is,kk'}$ represents the microbiome similarity between the sample $i$ and sample $s$ when sample $i$ is from study $k$ and the sample $s$ is from the study $k'$. Note that similarities $K_{is,kk'}, K_{jt,kk'}, K_{it,kk'}, K_{js,kk'}$ are the underlying similarities between subjects from different studies, which may not be available.

Note that, under the null hypothesis, $\tau = 0$, and the corresponding variance-covariance reduces. For example, $\mathrm{Cov}(Z_{ij,k}, Z_{st,k}|\mathbf{X},\mathbf{M}) = 0$ under the null, indicating that the trait similarity between pairs of samples in study $k$ is uncorrelated to the trait similarity between pairs of samples in study $k'$.

We can combine the mean relationship (A.1) and the variance-covariance (A.2), (A.3), (A.4), (A.5) of the trait similarities to construct SMRmix. Clearly, from (A.1), an intercept is needed.

Comparing (A.4) to (A.5), if pairs of samples come from the same study, an additional covariance term arises under the null. Thus, we would need an extra random effect in the SMRmix to capture the study effect.

Comparing (A.3) to (A.4), $(Z_{ij,k},\ Z_{it,k})$ has higher covariance than $(Z_{ij,k}, Z_{st,k})$ because they share the $i$-th sample. Thus, we need an extra random effect to capture this additional correlation.

Based on (A.2), a random error is needed. Thus, the proposed SMRmix model can be formulated as follows,

$$Z_{ij,k} = a + bK_{ij,k} + u_k + f_i + e_{ij,k},\ i < j,$$

where $a$ is an intercept, $b$ is the regression coefficient, $u_k$ is the study-specific random effect, $f_i$ is the individual-level random effect, and $e_{ij,k}$ is the pairwise similarity random effect.

Similarly, for binary outcomes, the same SMRmix model can be justified by comparing the following conditional variance and covariances (See details in section B.2),

$$\begin{aligned}\mathrm{Var}(Z_{ij,k}|\mathbf{X},\mathbf{M}) =& [\mu_{ik}^0(1-\mu_{ik}^0)\mu_{jk}^0(1-\mu_{jk}^0)]^2 \\ & \cdot [2\sigma^4 + \tau\sigma^2(K_{ii,k} + 2K_{ij,k} + K_{jj,k}) + \tau^2(K_{ii,k}K_{jj,k} + K_{ij,k}^2)] \\ & + [\mu_{ik}^0(1-\mu_{ik}^0)\mu_{jk}^0(1-\mu_{jk}^0)][\mu_{ik}^0(1-\mu_{ik}^0)(\sigma^2 + \tau K_{ii,k}) + \mu_{jk}^0(1-\mu_{jk}^0)(\sigma^2 + \tau K_{jj,k})] \\ & + \mu_{ik}^0(1-\mu_{ik}^0)\mu_{jk}^0(1-\mu_{jk}^0)[(1-2\mu_{ik}^0)(1-2\mu_{jk}^0)(\sigma^2 + \tau K_{ij,k}) + 1]\end{aligned} \tag{A.6}$$

$$\begin{aligned}\mathrm{Cov}(Z_{ij,k}, Z_{it,k}|\mathbf{X},\mathbf{M}) =& [\mu_{ik}^0(1-\mu_{ik}^0)]^2\mu_{jk}^0(1-\mu_{jk}^0)\mu_{tk}^0(1-\mu_{tk}^0) \\ & \cdot [2\sigma^4 + \tau\sigma^2(K_{it,k} + K_{ii,k} + K_{ji,k} + K_{jt,k}) + \tau^2(K_{ii,k}K_{jt,k} + K_{it,k}K_{ij,k})] \\ & + \mu_{ik}^0(1-\mu_{ik}^0)\mu_{jk}^0(1-\mu_{jk}^0)\mu_{tk}^0(1-\mu_{tk}^0)(\sigma^2 + \tau K_{jt,k})\end{aligned} \tag{A.7}$$

$$\begin{aligned}\mathrm{Cov}(Z_{ij,k}, Z_{st,k}|\mathbf{X},\mathbf{M}) =& \mu_{ik}^0(1-\mu_{ik}^0)\mu_{jk}^0(1-\mu_{jk}^0)\mu_{sk}^0(1-\mu_{sk}^0)\mu_{tk}^0(1-\mu_{tk}^0) \\ & \cdot [2\sigma^4 + \tau\sigma^2(K_{it,k} + K_{is,k} + K_{js,k} + K_{jt,k}) + \tau^2(K_{is,k}K_{jt,k} + K_{it,k}K_{js,k})]\end{aligned} \tag{A.8}$$

$$\begin{aligned}\mathrm{Cov}(Z_{ij,k}, Z_{st,k'}|\mathbf{X},\mathbf{M}) =& \mu_{ik}^0(1-\mu_{ik}^0)\mu_{jk}^0(1-\mu_{jk}^0)\mu_{sk'}^0(1-\mu_{sk'}^0)\mu_{tk'}^0(1-\mu_{tk'}^0) \\ & \cdot [\tau^2(K_{is,kk'}K_{jt,kk'} + K_{it,kk'}K_{js,kk'})]\end{aligned} \tag{A.9}$$

### B Conditional Variance-covariance Matrix of Z

In this section, we present the expressions of variance-covariance matrix $\mathbf{V} = \mathrm{Var}(\mathbf{Z}|\mathbf{X}, \mathbf{M})$ for both continuous case and binary case.

#### B.1 Continuous outcomes

For continuous outcomes, we assume a linear mixed-effect model,

$$Y_{ik} = \mathbf{X}_{ik}^T \beta + f(\mathbf{M}_{ik}) + h_k + \varepsilon_{ik},$$

where $h_k \sim N(0, \sigma^2)$, $\varepsilon_{ik} \sim N(0, \delta_k^2)$, and $f(\mathbf{M}_{ik})$ depicts the microbiome effect on the outcome. Let $\gamma_{ik} = f(\mathbf{M}_{ik})$, $\gamma_k = (\gamma_{1k}, \cdots, \gamma_{n_k k}) \sim \mathrm{MVN}(\mathbf{0}, \tau \mathbf{K}_k)$.

In similarity matrix regression mixed effect model (SMRmix), the trait similarity between $i$-th subject and $j$-th subject in $k$-th study is

$$Z_{ij,k} = (Y_{ik} - \hat{\mu}_{ik}^0)(Y_{jk} - \hat{\mu}_{jk}^0),$$

where $\hat{\mu}_{ik}^0$ is the fitted values of $Y_{ik}$ under the null hypothesis. Let $\gamma_{ik} = f(\mathbf{M}_{ik})$. The conditional expectation of $Z_{ij,k}$ $\mathbb{E}(Z_{ij,k}|\mathbf{X}, \mathbf{M})$ is

$$\begin{aligned}
&\mathbb{E}(Z_{ij,k}|\mathbf{X}, \mathbf{M}) = \mathbb{E}[(Y_{ik} - \mu_{ik}^0)(Y_{jk} - \mu_{jk}^0)|\mathbf{X}, \mathbf{M}] \\
=&\mathbb{E}[(\gamma_{ik} + h_k + \varepsilon_{ik})(\gamma_{jk} + h_k + \varepsilon_{jk})|\mathbf{X}, \mathbf{M}] \\
=&\sigma^2 + \tau K_{ij,k}.
\end{aligned}$$

By the law of total variance, the variance $\mathrm{Var}(Z_{ij,k}|\mathbf{X}, \mathbf{M})$ is

$$\mathrm{Var}(Z_{ij,k}|\mathbf{X}, \mathbf{M}) = \mathrm{Var}[\mathbb{E}(Z_{ij,k}|\mathbf{X}, \mathbf{M}, \gamma)|\mathbf{X}, \mathbf{M}] + \mathbb{E}[\mathrm{Var}(Z_{ij,k}|\mathbf{X}, \mathbf{M}, \gamma)|\mathbf{X}, \mathbf{M}],$$

where

$$
\begin{aligned}
\mathrm{Var}(Z_{ij,k}|\mathbf{X},\mathbf{M},\gamma) &= \mathrm{Var}((\gamma_{ik}+h_k+\varepsilon_{ik})(\gamma_{jk}+h_k+\varepsilon_{jk})|\mathbf{X},\mathbf{M},\gamma)\\
=&\mathrm{Var}((\gamma_{ik}+\gamma_{jk}+\varepsilon_{ik}+\varepsilon_{jk})h_k+\gamma_{ik}\varepsilon_{jk}+\gamma_{jk}\varepsilon_{ik}+h_k^2+\varepsilon_{ik}\varepsilon_{jk}|\mathbf{X},\mathbf{M},\gamma)\\
=&(\gamma_{ik}+\gamma_{jk})^2\sigma^2+(\gamma_{ik}^2+\gamma_{jk}^2)\delta_k^2+2\sigma^4+2\delta_k^2\sigma^2+\delta_k^4,
\end{aligned}
$$

and

$$
\mathbb{E}(Z_{ij,k}|\mathbf{X},\mathbf{M},\gamma) = \gamma_{ik}\gamma_{jk}+\sigma^2.
$$

Therefore,

$$
\begin{aligned}
\mathrm{Var}(Z_{ij,k}|\mathbf{X},\mathbf{M}) =&\tau\sigma^2(K_{ii,k}+2K_{ij,k}+K_{jj,k})+\tau^2(K_{ii,k}K_{jj,k}+K_{ij,k}^2)\\
&+2\sigma^4+\tau\delta_k^2(K_{ii,k}+K_{jj,k})+2\delta_k^2\sigma^2+\delta_k^4
\end{aligned}
$$

By the law of total covariance, the conditional covariance $\mathrm{Cov}(Z_{ij,k},Z_{it,k}|\mathbf{X},\mathbf{M})$ is

$$
\begin{aligned}
&\mathrm{Cov}(Z_{ij,k},Z_{it,k}|\mathbf{X},\mathbf{M})\\
=&\mathbb{E}[\mathrm{Cov}(Z_{ij,k},Z_{it,k}|\mathbf{X},\mathbf{M},\gamma)|\mathbf{X},\mathbf{M}]+\mathrm{Cov}[\mathbb{E}(Z_{ij,k}|\mathbf{X},\mathbf{M},\gamma),\mathbb{E}(Z_{it,k}|\mathbf{X},\mathbf{M},\gamma)|\mathbf{X},\mathbf{M}]
\end{aligned}
$$

where

$$
\begin{aligned}
\mathbb{E}[\mathrm{Cov}(Z_{ij,k},Z_{it,k}|\mathbf{X},\mathbf{M},\gamma)|\mathbf{X},\mathbf{M}] &= \mathbb{E}[(\gamma_{ik}+\gamma_{jk})(\gamma_{ik}+\gamma_{tk})\sigma^2+\gamma_{jk}\gamma_{tk}+\delta_k^2\sigma^2+2\sigma^4|\mathbf{X},\mathbf{M},\gamma]\\
=&\tau\sigma^2(K_{it,k}+K_{ii,k}+K_{ji,k}+K_{jt,k})+\tau\delta_k^2K_{jt,k}+\delta_k^2\sigma^2+2\sigma^4,
\end{aligned}
$$

and

$$\mathrm{Cov}[\mathbb{E}(Z_{ij,k}|\mathbf{X},\mathbf{M},\gamma),\mathbb{E}(Z_{it,k}|\mathbf{X},\mathbf{M},\gamma)|\mathbf{X},\mathbf{M}] = \mathrm{Cov}(\gamma_{ik}\gamma_{jk}+\sigma^2,\gamma_{ik}\gamma_{tk}+\sigma^2)$$
$$=\tau^2(K_{ii,k}K_{jt,k}+K_{it,k}K_{ij,k})$$

Therefore,

$$\begin{aligned}\mathrm{Cov}(Z_{ij,k},Z_{it,k}|\mathbf{X},\mathbf{M}) =& \tau\sigma^2(K_{it,k}+K_{ii,k}+K_{ji,k}+K_{jt,k})+\tau^2(K_{ii,k}K_{jt,k}+K_{it,k}K_{ij,k})\\ &+2\sigma^4+\tau\delta_k^2K_{jt,k}+\delta_k^2\sigma^2\end{aligned}$$

By the law of total covariance, the conditional covariance $\mathrm{Cov}(Z_{ij,k},Z_{st,k}|\mathbf{X},\mathbf{M})$ is

$$\begin{aligned}&\mathrm{Cov}(Z_{ij,k},Z_{it,k}|\mathbf{X},\mathbf{M})\\ =&\mathbb{E}[\mathrm{Cov}(Z_{ij,k},Z_{st,k}|\mathbf{X},\mathbf{M},\gamma)|\mathbf{X},\mathbf{M}]+\mathrm{Cov}[\mathbb{E}(Z_{ij,k}|\mathbf{X},\mathbf{M},\gamma),\mathbb{E}(Z_{st,k}|\mathbf{X},\mathbf{M},\gamma)|\mathbf{X},\mathbf{M}]\end{aligned}$$

where

$$\mathbb{E}[\mathrm{Cov}(Z_{ij,k},Z_{st,k}|\mathbf{X},\mathbf{M},\gamma)|\mathbf{X},\mathbf{M}] = \mathbb{E}[(\gamma_{ik}+\gamma_{jk})(\gamma_{sk}+\gamma_{tk})\sigma^2+2\sigma^4|\mathbf{X},\mathbf{M}]$$
$$=\tau\sigma^2(K_{it,k}+K_{is,k}+K_{js,k}+K_{jt,k})+2\sigma^4,$$

and

$$\mathrm{Cov}[\mathbb{E}(Z_{ij,k}|\mathbf{X},\mathbf{M},\gamma),\mathbb{E}(Z_{st,k}|\mathbf{X},\mathbf{M},\gamma)|\mathbf{X},\mathbf{M}] = \mathrm{Cov}(\gamma_{ik}\gamma_{jk}+\sigma^2,\gamma_{sk}\gamma_{tk}+\sigma^2|\mathbf{X},\mathbf{M})$$
$$=\tau^2(K_{is,k}K_{jt,k}+K_{it,k}K_{js,k})$$

Therefore,

$$\mathrm{Cov}(Z_{ij,k}, Z_{st,k}|\mathbf{X},\mathbf{M}) = \tau\sigma^2(K_{it,k} + K_{is,k} + K_{js,k} + K_{jt,k}) + \tau^2(K_{is,k}K_{jt,k} + K_{it,k}K_{js,k}) + 2\sigma^4$$

By the law of total covariance, the conditional covariance $\mathrm{Cov}(Z_{ij,k}, Z_{st,k'}|\mathbf{X},\mathbf{M})$ is

$$\begin{aligned}
&\mathrm{Cov}(Z_{ij,k}, Z_{st,k'}|\mathbf{X},\mathbf{M}) \\
=&\mathbb{E}[\mathrm{Cov}(Z_{ij,k}, Z_{st,k'}|\mathbf{X},\mathbf{M},\gamma)|\mathbf{X},\mathbf{M}] + \mathrm{Cov}[\mathbb{E}(Z_{ij,k}|\mathbf{X},\mathbf{M},\gamma), \mathbb{E}(Z_{st,k'}|\mathbf{X},\mathbf{M},\gamma)|\mathbf{X},\mathbf{M}] \\
=&\tau^2(K_{is,kk'}K_{jt,kk'} + K_{it,kk'}K_{js,kk'})
\end{aligned}$$

where we assume $K_{is,kk'}$ is a known value of similarity between $i$-th subject from $k$-th study and $s$-th subject from $k'$-th study.

#### B.2 Binary outcomes

For binary outcomes, the mixed-effect logistic model is

$$g(\mathbb{P}(Y_{ik} = 1)) = \mathbf{X}_{ik}^T\beta + f(\mathbf{M}_{ik}) + h_k,$$

where $g(\cdot)$ is the logit function, $h_k \sim N(0,\sigma^2)$, and $f(\mathbf{M}_{ik})$ depicts the microbiome effect on the outcome. Let $\gamma_{ik} = f(\mathbf{M}_{ik})$, $\gamma_k = (\gamma_{1k}, \cdots, \gamma_{n_k k}) \sim \mathrm{MVN}(\mathbf{0}, \tau\mathbf{K}_k)$.

Let $\mu_{ik}(\gamma_{ik}, h_k) = \mathbb{E}(Y_{ik}|\mathbf{X},\mathbf{M},\gamma,\mathbf{h}) = g^{-1}(\mathbf{X}_{ik}^T\beta + \gamma_{ik} + h_k)$, and $\mu_{ik}^0 = \mu_{ik}(0,0) = g^{-1}(\mathbf{X}_{ik}^T\beta)$.

By Taylor expansion of $\mu_{ik}(\gamma_{ik}, h_k)$ around $(0,0)$, the conditional expectation of $Z_{ij,k}$ is,

$$
\begin{aligned}
&\mathbb{E}(Z_{ij,k}|\mathbf{X},\mathbf{M}) = \mathbb{E}((Y_{ik}-\mu^0_{ik})(Y_{jk}-\mu^0_{jk})|\mathbf{X},\mathbf{M}) \\
=&\mathbb{E}\{\mathbb{E}[(Y_{ik}-\mu_{ik}(\gamma_{ik},h_k)+\mu_{ik}(\gamma_{ik},h_k)-\mu^0_{ik})(Y_{jk}-\mu_{jk}(\gamma_{jk},h_k)+\mu_{jk}(\gamma_{jk},h_k)-\mu^0_{jk})|\mathbf{X},\mathbf{M},\gamma,\mathbf{h}]|\mathbf{X},\mathbf{M}\} \\
=&\mathbb{E}[(\mu_{ik}(\gamma_{ik},h_k)-\mu^0_{ik})(\mu_{jk}(\gamma_{jk},h_k)-\mu^0_{jk})|\mathbf{X},\mathbf{M}] \\
\approx&\mathbb{E}[(g^{-1})'(\mathbf{X}^T_{ik}\beta)(\gamma_{ik}+h_k)(g^{-1})'(\mathbf{X}^T_{jk}\beta)(\gamma_{jk}+h_k)|\mathbf{X},\mathbf{M}] \\
=&\mu^0_{ik}(1-\mu^0_{ik})\mu^0_{jk}(1-\mu^0_{jk})(\sigma^2+\tau K_{ij,k})
\end{aligned}
$$

By the law of total variance, the variance $\mathrm{Var}(Z_{ij,k}|\mathbf{X},\mathbf{M})$ is

$$
\mathrm{Var}(Z_{ij,k}|\mathbf{X},\mathbf{M}) = \mathrm{Var}[\mathbb{E}(Z_{ij,k}|\mathbf{X},\mathbf{M},\gamma,\mathbf{h})|\mathbf{X},\mathbf{M}] + \mathbb{E}[\mathrm{Var}(Z_{ij,k}|\mathbf{X},\mathbf{M},\gamma,\mathbf{h})|\mathbf{X},\mathbf{M}],
$$

where

$$
\begin{aligned}
&\mathrm{Var}[\mathbb{E}(Z_{ij,k}|\mathbf{X},\mathbf{M},\gamma,\mathbf{h})|\mathbf{X},\mathbf{M}] \approx \mathrm{Var}[(g^{-1})'(\mathbf{X}^T_{ik}\beta)(\gamma_{ik}+h_k)(g^{-1})'(\mathbf{X}^T_{jk}\beta)(\gamma_{jk}+h_k)|\mathbf{X},\mathbf{M}] \\
&= [\mu^0_{ik}(1-\mu^0_{ik})\mu^0_{jk}(1-\mu^0_{jk})]^2[2\sigma^4+\tau\sigma^2(K_{ii,k}+2K_{ij,k}+K_{jj,k})+\tau^2(K_{ii,k}K_{jj,k}+K^2_{ij,k})], 2qw
\end{aligned}
$$

and

$$
\begin{aligned}
&\mathbb{E}[\mathrm{Var}(Z_{ij,k}|\mathbf{X},\mathbf{M},\gamma,\mathbf{h})|\mathbf{X},\mathbf{M}] \\
=&\mathbb{E}\{\mathrm{Var}[(Y_{ik}-\mu_{ik}(\gamma_{ik},h_k)+\mu_{ik}(\gamma_{ik},h_k)-\mu^0_{ik})(Y_{jk}-\mu_{jk}(\gamma_{jk},h_k)+\mu_{jk}(\gamma_{jk},h_k)-\mu^0_{jk})|\mathbf{X},\mathbf{M},\gamma,\mathbf{h}]|\mathbf{X},\mathbf{M}\} \\
\approx&\mathbb{E}\{\mathrm{Var}[(Y_{ik}-\mu_{ik}(\gamma_{ik},h_k))(Y_{jk}-\mu_{jk}(\gamma_{jk},h_k))|\mathbf{X},\mathbf{M},\gamma,\mathbf{h}] \\
&+(\mu_{jk}(\gamma_{jk},h_k)-\mu^0_{jk})(Y_{ik}-\mu_{ik}(\gamma_{ik},h_k))+(\mu_{ik}(\gamma_{ik},h_k)-\mu^0_{ik})(Y_{jk}-\mu_{jk}(\gamma_{jk},h_k))|\mathbf{X},\mathbf{M}\} \\
=&\mathbb{E}[\mu_{ik}(\gamma_{ik},h_k)(1-\mu_{ik}(\gamma_{ik},h_k))\mu_{jk}(\gamma_{jk},h_k)(1-\mu_{jk}(\gamma_{jk},h_k)) \\
&+(\mu_{ik}(\gamma_{ik},h_k)-\mu^0_{ik})^2\mu_{jk}(\gamma_{jk},h_k)(1-\mu_{jk}(\gamma_{jk},h_k)) \\
&+(\mu_{jk}(\gamma_{jk},h_k)-\mu^0_{jk})^2\mu_{ik}(\gamma_{ik},h_k)(1-\mu_{ik}(\gamma_{ik},h_k))|\mathbf{X},\mathbf{M}]
\end{aligned}
$$

$$\approx \mathbb{E}\{(1-2\mu_{ik}^0)(g^{-1})'(\mathbf{X}_{ik}^T\beta)(\gamma_{ik}+h_k)(1-2\mu_{jk}^0)(g^{-1})'(\mathbf{X}_{jk}^T\beta)(\gamma_{jk}+h_k)|\mathbf{X},\mathbf{M}\}+\mu_{ik}^0(1-\mu_{ik}^0)\mu_{jk}^0(1-\mu_{jk}^0)$$
$$+\mu_{jk}^0(1-\mu_{jk}^0)[(g^{-1})'(\mathbf{X}_{ik}^T\beta)]^2(\sigma^2+\tau K_{ii,k})+\mu_{ik}^0(1-\mu_{ik}^0)[(g^{-1})'(\mathbf{X}_{jk}^T\beta)]^2(\sigma^2+\tau K_{jj,k})$$
$$=\mu_{ik}^0(1-\mu_{ik}^0)\mu_{jk}^0(1-\mu_{jk}^0)[(1-2\mu_{ik}^0)(1-2\mu_{jk}^0)(\sigma^2+\tau K_{ij,k})+1]$$
$$+[\mu_{ik}^0(1-\mu_{ik}^0)\mu_{jk}^0(1-\mu_{jk}^0)][\mu_{ik}^0(1-\mu_{ik}^0)(\sigma^2+\tau K_{ii,k})+\mu_{jk}^0(1-\mu_{jk}^0)(\sigma^2+\tau K_{jj,k})].$$

Therefore,

$$\begin{aligned}\mathrm{Var}(Z_{ij,k}|\mathbf{X},\mathbf{M}) =&[\mu_{ik}^0(1-\mu_{ik}^0)\mu_{jk}^0(1-\mu_{jk}^0)]^2[2\sigma^4+\tau\sigma^2(K_{ii,k}+2K_{ij,k}+K_{jj,k})+\tau^2(K_{ii,k}K_{jj,k}+K_{ij,k}^2)]\\ &+\mu_{ik}^0(1-\mu_{ik}^0)\mu_{jk}^0(1-\mu_{jk}^0)[(1-2\mu_{ik}^0)(1-2\mu_{jk}^0)(\sigma^2+\tau K_{ij,k})+1]\\ &+[\mu_{ik}^0(1-\mu_{ik}^0)\mu_{jk}^0(1-\mu_{jk}^0)][\mu_{ik}^0(1-\mu_{ik}^0)(\sigma^2+\tau K_{ii,k})+\mu_{jk}^0(1-\mu_{jk}^0)(\sigma^2+\tau K_{jj,k})].\end{aligned}$$

By the law of total covariance, the conditional covariance $\mathrm{Cov}(Z_{ij,k},Z_{it,k}|\mathbf{X},\mathbf{M})$ is

$$\begin{aligned}&\mathrm{Cov}(Z_{ij,k},Z_{it,k}|\mathbf{X},\mathbf{M})\\ =&\mathbb{E}[\mathrm{Cov}(Z_{ij,k},Z_{it,k}|\mathbf{X},\mathbf{M},\gamma,\mathbf{h})|\mathbf{X},\mathbf{M}]+\mathrm{Cov}[\mathbb{E}(Z_{ij,k}|\mathbf{X},\mathbf{M},\gamma,\mathbf{h}),\mathbb{E}(Z_{it,k}|\mathbf{X},\mathbf{M},\gamma,\mathbf{h})|\mathbf{X},\mathbf{M}]\end{aligned}$$

where

$$\begin{aligned}&\mathbb{E}[\mathrm{Cov}(Z_{ij,k},Z_{it,k}|\mathbf{X},\mathbf{M},\gamma,\mathbf{h})|\mathbf{X},\mathbf{M}]\\ =&\ \mathbb{E}[\mathrm{Cov}((Y_{ik}-\mu_{ik}(\gamma_{ik},h_k)+\mu_{ik}(\gamma_{ik},h_k)-\mu_{ik}^0)(Y_{jk}-\mu_{jk}(\gamma_{jk},h_k)+\mu_{jk}(\gamma_{jk},h_k)-\mu_{jk}^0),\\ &\qquad (Y_{ik}-\mu_{ik}(\gamma_{ik},h_k)+\mu_{ik}(\gamma_{ik},h_k)-\mu_{ik}^0)(Y_{tk}-\mu_{tk}(\gamma_{tk},h_k)+\mu_{tk}(\gamma_{tk},h_k)-\mu_{tk}^0)|\mathbf{X},\mathbf{M},\gamma,\mathbf{h})|\mathbf{X},\mathbf{M}]\\ =&\ \mathbb{E}[(\mu_{jk}-\mu_{jk}^0)(\mu_{tk}-\mu_{tk}^0)\mu_{ik}(\gamma_{ik},h_k)(1-\mu_{ik}(\gamma_{ik},h_k))|\mathbf{X},\mathbf{M}]\\ \approx&\ (g^{-1})'(\mu_{jk}^0)(g^{-1})'(\mu_{tk}^0)\mathbb{E}[(\gamma_{jk}+h_k)(\gamma_{tk}+h_k)(\mu_{ik}^0(1-\mu_{ik}^0)+(1-2\mu_{ik}^0)(g^{-1})'(\mu_{ik}^0)(\gamma_{ik}+h_k))]\\ =&\ \mu_{ik}^0(1-\mu_{ik}^0)\mu_{jk}^0(1-\mu_{jk}^0)\mu_{tk}^0(1-\mu_{tk}^0)(\sigma^2+\tau K_{jt,k}),\end{aligned}$$

and

$$
\begin{aligned}
&\mathrm{Cov}[\mathbb{E}(Z_{ij,k}|\mathbf{X},\mathbf{M},\gamma,\mathbf{h}),\mathbb{E}(Z_{it,k}|\mathbf{X},\mathbf{M},\gamma,\mathbf{h})|\mathbf{X},\mathbf{M}] \\
\approx&\mathrm{Cov}((g^{-1})'(\mathbf{X}_{ik}^T\beta)(g^{-1})'(\mathbf{X}_{jk}^T\beta)(\gamma_{ik}+h_k)(\gamma_{jk}+h_k),(g^{-1})'(\mathbf{X}_{ik}^T\beta)(g^{-1})'(\mathbf{X}_{tk}^T\beta)(\gamma_{ik}+h_k)(\gamma_{tk}+h_k)|\mathbf{X},\mathbf{M}) \\
=&[\mu_{ik}^0(1-\mu_{ik}^0)]^2\mu_{jk}^0(1-\mu_{jk}^0)\mu_{tk}^0(1-\mu_{tk}^0)[\tau^2(K_{ii,k}K_{jt,k}+K_{it,k}K_{ij,k})+\tau\sigma^2(K_{it,k}+K_{ii,k}+K_{ji,k}+K_{jt,k})+2\sigma^4]
\end{aligned}
$$

Therefore,

$$
\begin{aligned}
&\mathrm{Cov}(Z_{ij,k},Z_{it,k}|\mathbf{X},\mathbf{M}) = \mu_{ik}^0(1-\mu_{ik}^0)\mu_{jk}^0(1-\mu_{jk}^0)\mu_{tk}^0(1-\mu_{tk}^0)(\sigma^2+\tau K_{jt,k}) \\
&+[\mu_{ik}^0(1-\mu_{ik}^0)]^2\mu_{jk}^0(1-\mu_{jk}^0)\mu_{tk}^0(1-\mu_{tk}^0)[\tau^2(K_{ii,k}K_{jt,k}+K_{it,k}K_{ij,k})+\tau\sigma^2(K_{it,k}+K_{ii,k}+K_{ji,k}+K_{jt,k})+2\sigma^4]
\end{aligned}
$$

By the law of total covariance and conditional independence, $\mathrm{Cov}(Z_{ij,k},Z_{st,k}|\mathbf{X},\mathbf{M})$ is

$$
\begin{aligned}
&\mathrm{Cov}(Z_{ij,k},Z_{st,k}|\mathbf{X},\mathbf{M}) = \mathrm{Cov}[\mathbb{E}(Z_{ij,k}|\mathbf{X},\mathbf{M},\gamma,\mathbf{h}),\mathbb{E}(Z_{st,k}|\mathbf{X},\mathbf{M},\gamma,\mathbf{h})|\mathbf{X},\mathbf{M}] \\
\approx&\mathrm{Cov}((g^{-1})'(\mathbf{X}_{ik}^T\beta)(g^{-1})'(\mathbf{X}_{jk}^T\beta)(\gamma_{ik}+h_k)(\gamma_{jk}+h_k),(g^{-1})'(\mathbf{X}_{sk}^T\beta)(g^{-1})'(\mathbf{X}_{tk}^T\beta)(\gamma_{sk}+h_k)(\gamma_{tk}+h_k)|\mathbf{X},\mathbf{M}) \\
=&\mu_{ik}^0(1-\mu_{ik}^0)\mu_{jk}^0(1-\mu_{jk}^0)\mu_{sk}^0(1-\mu_{sk}^0)\mu_{tk}^0(1-\mu_{tk}^0) \\
&\cdot[2\sigma^4+\tau\sigma^2(K_{it,k}+K_{is,k}+K_{js,k}+K_{jt,k})+\tau^2(K_{is,k}K_{jt,k}+K_{it,k}K_{js,k})]
\end{aligned}
$$

Similarly, by the law of total covariance and conditional independence, $\mathrm{Cov}(Z_{ij,k},Z_{st,k'}|\mathbf{X},\mathbf{M})$ is

$$
\begin{aligned}
&\mathrm{Cov}(Z_{ij,k},Z_{st,k'}|\mathbf{X},\mathbf{M}) = \mathrm{Cov}[\mathbb{E}(Z_{ij,k}|\mathbf{X},\mathbf{M},\gamma,\mathbf{h}),\mathbb{E}(Z_{st,k'}|\mathbf{X},\mathbf{M},\gamma,\mathbf{h})|\mathbf{X},\mathbf{M}] \\
\approx&\mathrm{Cov}((g^{-1})'(\mathbf{X}_{ik}^T\beta)(g^{-1})'(\mathbf{X}_{jk}^T\beta)(\gamma_{ik}+h_k)(\gamma_{jk}+h_k),(g^{-1})'(\mu_{sk'}^0)(g^{-1})'(\mu_{tk'}^0)(\gamma_{sk'}+h_{k'})(\gamma_{tk'}+h_{k'})|\mathbf{X},\mathbf{M}) \\
=&\mu_{ik}^0(1-\mu_{ik}^0)\mu_{jk}^0(1-\mu_{jk}^0)\mu_{sk'}^0(1-\mu_{sk'}^0)\mu_{tk'}^0(1-\mu_{tk'}^0)[\tau^2(K_{is,kk'}K_{jt,kk'}+K_{it,kk'}K_{js,kk'})]
\end{aligned}
$$

### References

Tzeng, J.-Y., Zhang, D., Chang, S.-M., Thomas, D. C., and Davidian, M. (2009). Gene-Trait Similarity Regression for Multimarker-Based Association Analysis. *Biometrics*, 65(3):822–832.

Zhan, X. (2019). Relationship Between MiRKAT and Coefficient of Determination in Similarity Matrix Regression. *Processes*, 7(2):79.

### C Supplementary Materials for Simulation Studies and Application

Table S1: Summary of patient demographics, including age, gender and BMI, of 11 CRC datasets.

| | **FengQ_2015** | | **GuptaA_2019** | | **HanniganGD_2017** | | **ThomasAM_2018a** | | **ThomasAM_2018b** | | **ThomasAM_2019_c** | |
|---|---|---|---|---|---|---|---|---|---|---|---|---|
| | **CRC (N=10)** | **Controls (N=6)** | **CRC (N=23)** | **Controls (N=29)** | **CRC (N=19)** | **Controls (N=23)** | **CRC (N=4)** | **Controls (N=5)** | **CRC (N=25)** | **Controls (N=20)** | **CRC (N=26)** | **Controls (N=19)** |
| **Age** | | | | | | | | | | | | |
| Mean (SD) | 54.1 (7.69) | 56.8 (9.79) | 56.8 (6.02) | 40.5 (16.2) | 53.8 (7.49) | 52.3 (7.06) | 62.0 (1.83) | 61.6 (2.30) | 55.7 (7.44) | 54.6 (7.03) | 52.6 (11.3) | 52.6 (8.85) |
| **BMI** | | | | | | | | | | | | |
| Mean (SD) | 25.2 (2.55) | 23.5 (2.69) | 20.5 (1.45) | 22.5 (3.74) | 31.9 (8.33) | 26.9 (5.31) | 25.0 (3.56) | 24.2 (3.03) | 26.7 (4.64) | 25.0 (3.85) | 22.6 (2.90) | 22.9 (2.74) |
| **Gender** | | | | | | | | | | | | |
| Female | 7 (70.0%) | 3 (50.0%) | 9 (39.1%) | 18 (62.1%) | 6 (31.6%) | 15 (65.2%) | 0 (0%) | 1 (20.0%) | 7 (28.0%) | 7 (35.0%) | 11 (42.3%) | 8 (42.1%) |
| Male | 3 (30.0%) | 3 (50.0%) | 14 (60.9%) | 11 (37.9%) | 13 (68.4%) | 8 (34.8%) | 4 (100%) | 4 (80.0%) | 18 (72.0%) | 13 (65.0%) | 15 (57.7%) | 11 (57.9%) |

| | **VogtmannE_2016** | | **WirbelJ_2018** | | **YachidaS_2019** | | **YuJ_2015** | | **ZellerG_2014** | |
|---|---|---|---|---|---|---|---|---|---|---|
| | **CRC (N=30)** | **Controls (N=33)** | **CRC (N=33)** | **Controls (N=53)** | **CRC (N=144)** | **Controls (N=135)** | **CRC (N=16)** | **Controls (N=31)** | **CRC (N=24)** | **Controls (N=39)** |
| **Age** | | | | | | | | | | |
| Mean (SD) | 53.2 (10.3) | 55.0 (8.47) | 54.5 (8.47) | 52.3 (10.2) | 56.2 (7.83) | 52.2 (9.50) | 58.6 (5.16) | 62.1 (2.93) | 57.4 (6.53) | 55.4 (10.5) |
| **BMI** | | | | | | | | | | |
| Mean (SD) | 24.9 (4.46) | 25.1 (3.76) | 26.9 (4.13) | 24.8 (3.19) | 23.3 (3.56) | 22.5 (3.39) | 24.6 (2.95) | 23.5 (3.33) | 24.9 (3.14) | 24.7 (3.60) |
| **Gender** | | | | | | | | | | |
| Female | 6 (20.0%) | 8 (24.2%) | 16 (48.5%) | 26 (49.1%) | 51 (35.4%) | 64 (47.4%) | 8 (50.0%) | 12 (38.7%) | 8 (33.3%) | 23 (59.0%) |
| Male | 24 (80.0%) | 25 (75.8%) | 17 (51.5%) | 27 (50.9%) | 93 (64.6%) | 71 (52.6%) | 8 (50.0%) | 19 (61.3%) | 16 (66.7%) | 16 (41.0%) |

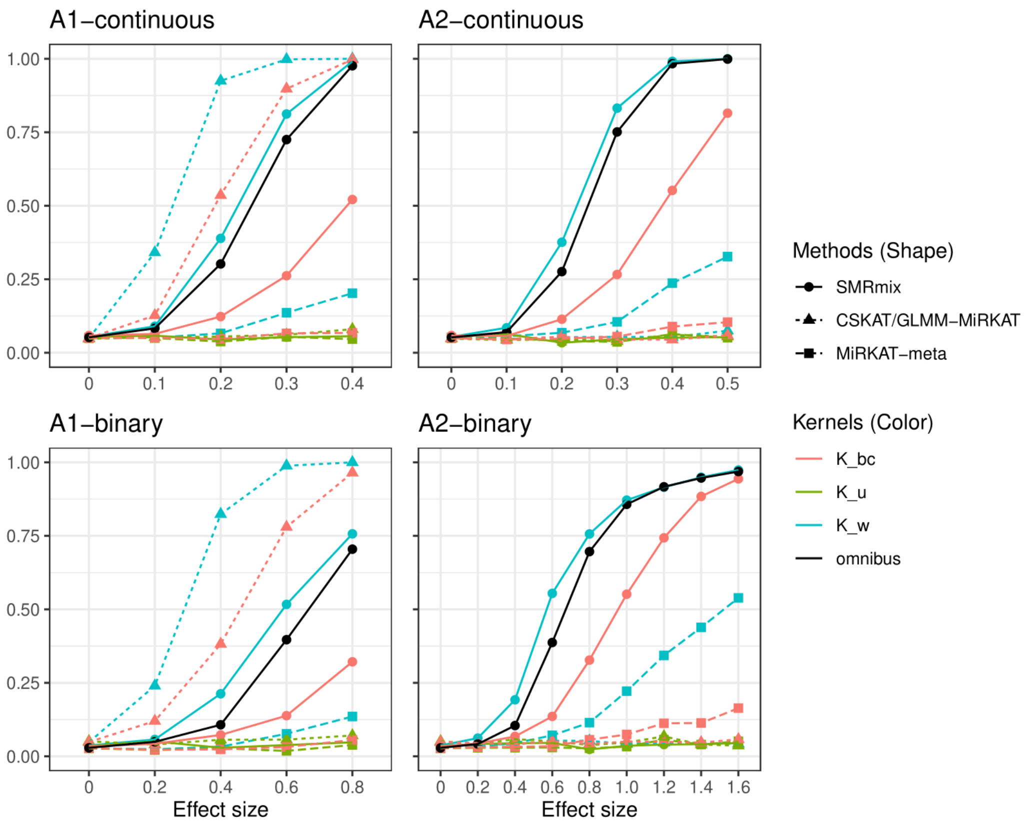


Figure S1: Type I Error and Power comparing CSKAT/GLMM-MiRKAT and SMRmix based on different kernels when differential biases exist. For continuous scenarios (upper two panels), $\delta_k^2 = 1, \forall k$, and $\sigma^2 = 1$.

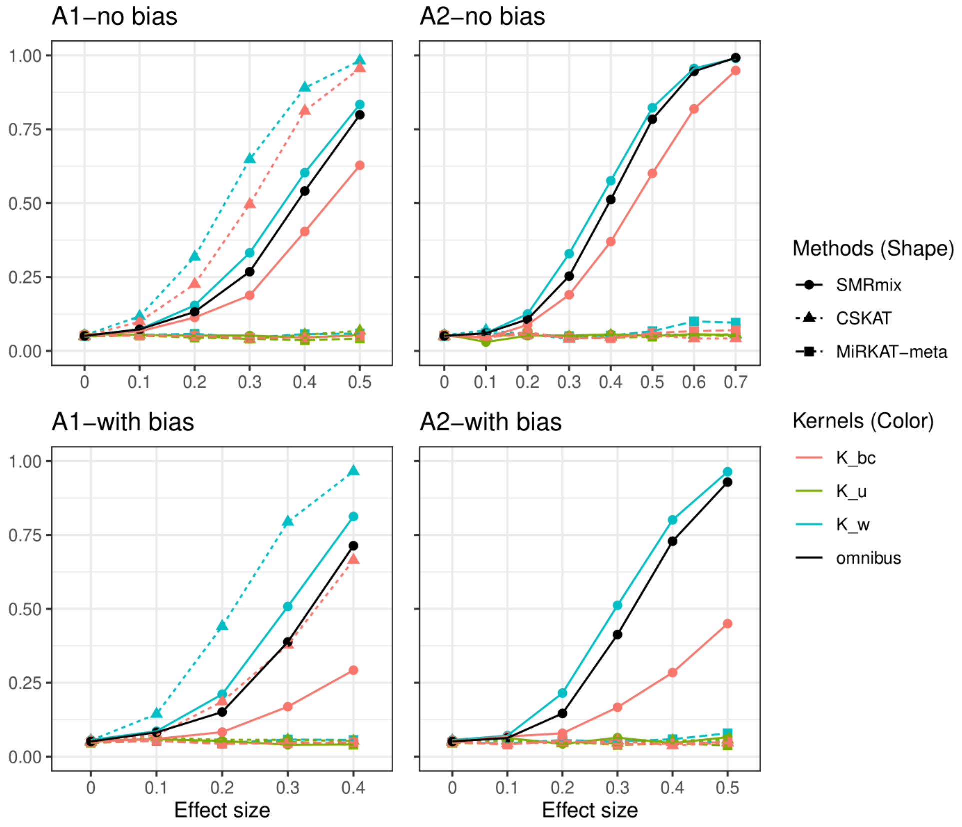


Figure S2: Type I Error and Power comparing CSKAT and SMRmix based on different kernels when the outcome is continuous with $\sigma^2 = 1$ and studies have different $\delta_k^2$. $\delta_k^2 = 1$, for $k = 1,2,3$, $\delta_k^2 = 3$, for $k = 4,5,6,7$, $\delta_k^2 = 5$, for $k = 8,9,10$.

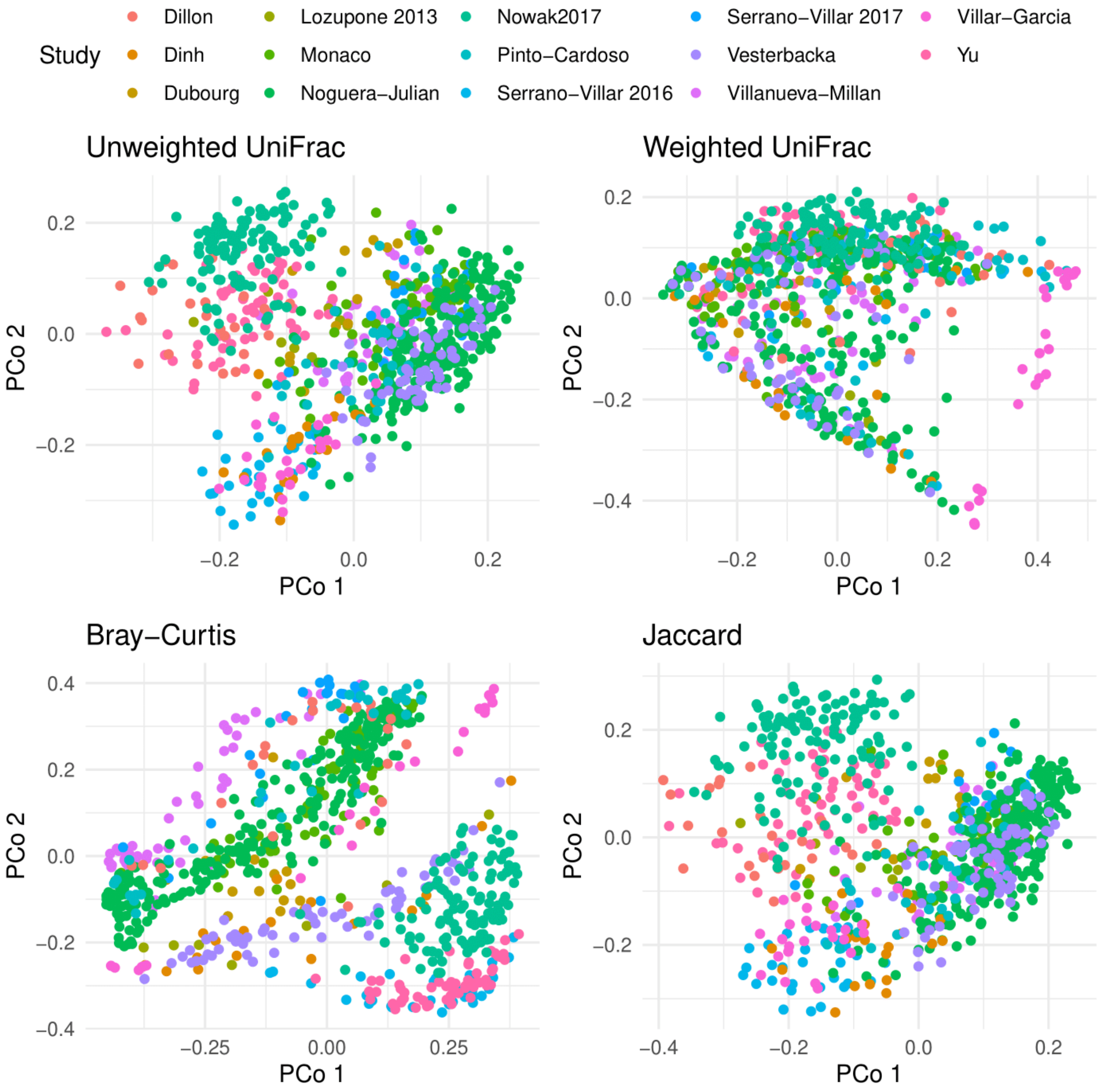


Figure S3: Visualization of Beta-diversities for 14 HIV Datasets

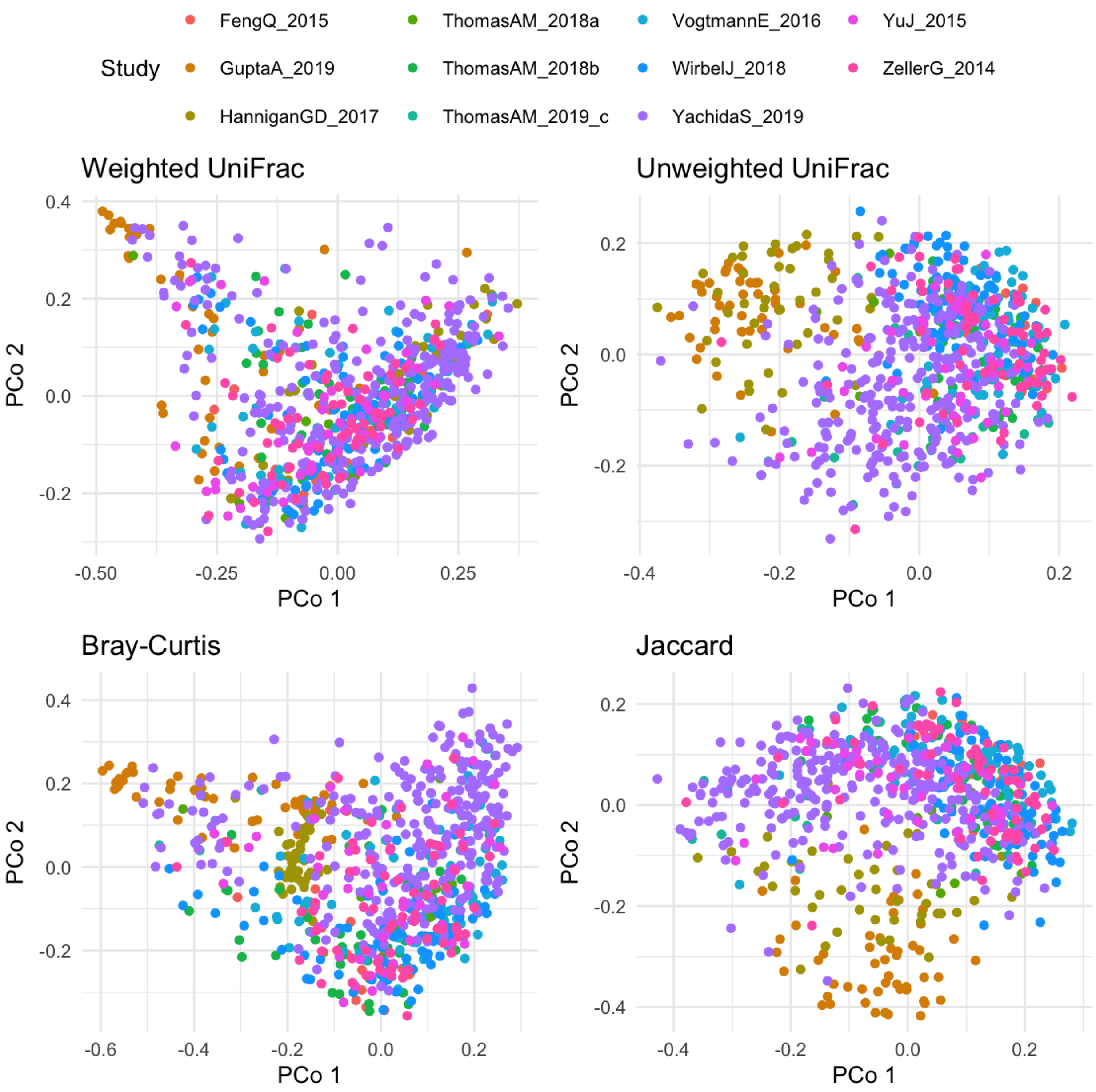


Figure S4: Visualization of Beta-diversities for 11 CRC Datasets
